# Ovalbumin antigen-specific activation of T cell receptor closely resembles soluble antibody stimulation as revealed by BOOST phosphotyrosine proteomics

**DOI:** 10.1101/2021.03.25.436968

**Authors:** Xien Yu Chua, Arthur Salomon

**Affiliations:** Department of Molecular Pharmacology, Physiology, and Biotechnology, Brown University, Providence, Rhode Island, USA; Department of Molecular Biology, Cell Biology and Biochemistry, Brown University, Providence, Rhode Island, USA

**Keywords:** T cell receptor, T cell signaling, BOOST, TMT, phosphotyrosine proteomics, pMHC, OT-1, OVA

## Abstract

Activation of T cell receptors (TCR) leads to a network of early signaling predominantly orchestrated by tyrosine phosphorylation in T cells. TCR are commonly activated using soluble anti-TCR antibodies, but this approach is not antigen-specific. Alternatively, activating the TCR using specific antigens of a range of binding affinities in the form of peptide-major histocompatibility complex (pMHC) is presumed to be more physiological. However, due to the lack of wide-scale phosphotyrosine (pTyr) proteomic studies directly comparing anti-TCR antibodies and pMHC, a comprehensive definition of these activated states remains enigmatic. Elucidation of the tyrosine phosphoproteome using quantitative pTyr proteomics enables a better understanding of the unique features of these activating agents and the role of ligand binding affinity on signaling. Here, we apply the recently established Broad-spectrum Optimization Of Selective Triggering (BOOST) to examine perturbations in tyrosine phosphorylation of TCR triggered by anti-TCR antibodies and pMHC. Our data reveals that high-affinity ovalbumin (OVA) pMHC activation of the TCR triggers a largely similar, albeit potentially stronger, pTyr-mediated signaling regulatory axis compared to anti-TCR antibody. Signaling output resulting from OVA pMHC variants correlates well with their weaker affinities, enabling affinity-tunable control of signaling strength. Collectively, we provide a framework for applying BOOST to compare pTyr-mediated signaling pathways of T cells activated in an antigen-independent and antigen-specific manner.

**Abstract Graphic:** 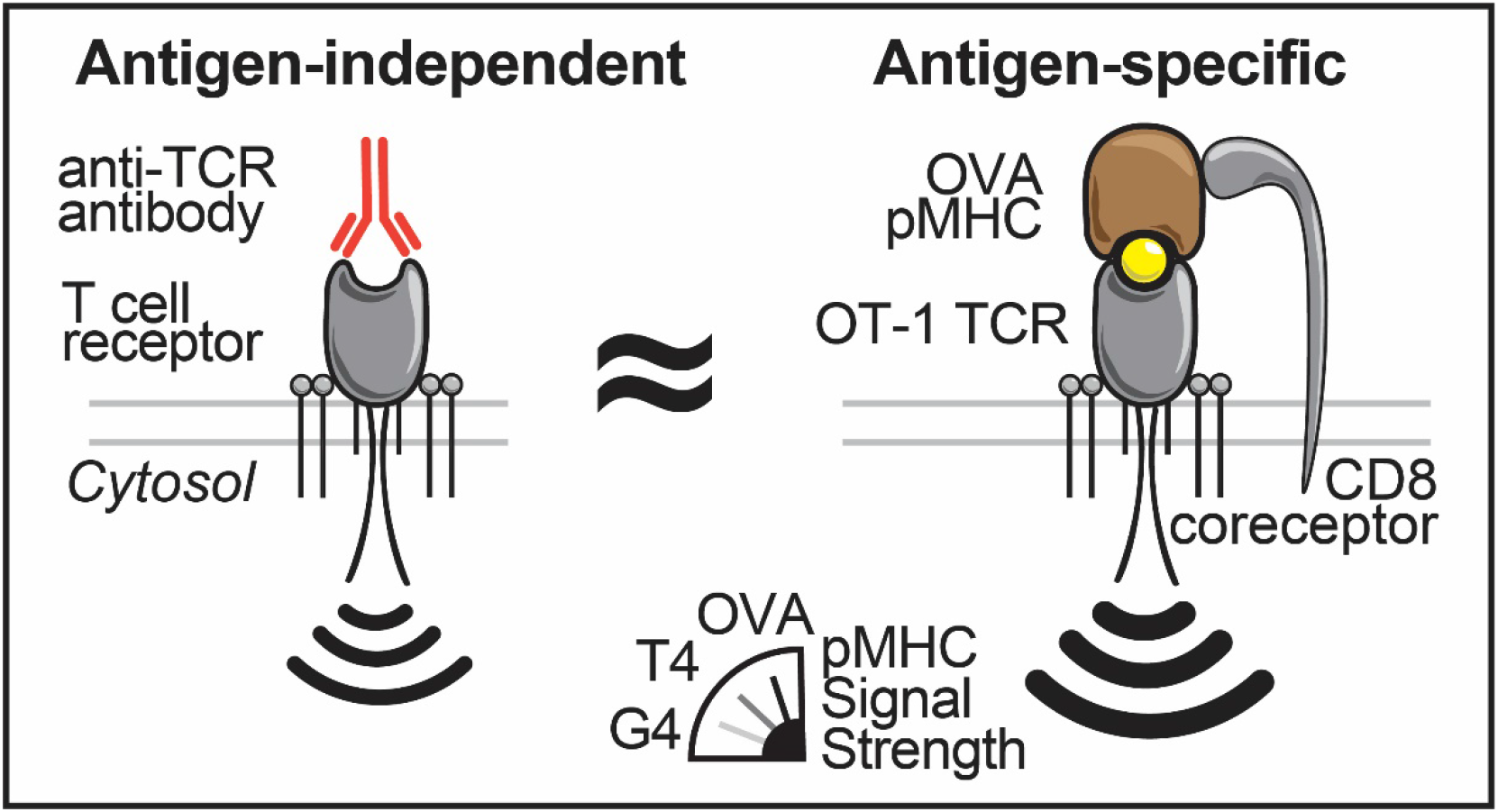

## Introduction

The interaction between TCR and its antigenic ligand is integral to how T cells regulate their physiological functions. Antigen-presenting cells present these antigens in the form of pMHC on their cell surface. The detection and binding of TCR to its cognate pMHC aided by a coreceptor stimulation enables an extracellular interaction to be transmitted across the plasma membrane as intracellular signaling events, leading to a battery of T cell responses. Early TCR signaling is predominantly driven by tyrosine phosphorylation, a post-translational modification (PTM). Upon the engagement of TCR to pMHC, the Src family kinase Lck phosphorylates immunoreceptor tyrosine-based activation motifs (ITAM) within CD3 and ζ chains of the TCR complex^1^. Each ITAM consist of a pair of tyrosine phosphorylation sites spaced by a highly conserved sequence motif with a defined interval^2^. ITAM phosphorylation provides the docking sites necessary to nucleate the formation of signaling complexes. This enables the propagation of signaling events that is required for cellular processes such as T cell differentiation and proliferation. However, early signaling events mediated by an array of tyrosine phosphorylation is incompletely understood due to the challenges in studying low-abundance tyrosine phosphorylation in the proteome (<1% of total phosphorylation^3^) and the limitations of methods using phosphorylation-specific antibodies^4, 5^.

Mass spectrometry(MS)-based pTyr proteomics has become an attractive method to examine the perturbations of tyrosine phosphorylation across the proteome in a high-throughput and unbiased manner. To overcome the challenges in identifying and quantifying low-abundance tyrosine phosphorylation in mass spectrometry, we recently developed the BOOST approach to increase the quantitation depth of tyrosine phosphoproteome while maintaining quantitative accuracy^6^. Briefly, pervanadate (PV) boost channels were introduced in a TMT experiment to trigger the selective fragmentation of pTyr peptides, facilitating the quantitation of reporter ions in non-boost channels in a data-dependent acquisition (DDA) mode. This is possible because PV is a potent broad-spectrum tyrosine phosphatase inhibitor that elevates the abundance of pTyr sites across the proteome^7^, thereby increasing the intensity of multiplexed pTyr-containing precursor ions. However, it has not been demonstrated that BOOST can be applied beyond a proof-of-concept study to gain useful biological insights in the immune system. Here, we attempt to reveal the differences in pTyr-mediated signaling pathways, or the lack thereof, in antigen-independent versus antigen-specific activation of TCR.

Due to the ease of use, many proteomic studies investigating T cell signaling pathways have commonly relied on antigen-independent antibody-based activation of T cells^8, 9^. Monoclonal antibodies such as C305^10^ and OKT3^11^ are popular reagents used to activate T cells by binding to the TCR to mimic the clustering of TCR when TCR engages with a pMHC. However, TCR triggering using antibodies is not antigen-specific^12^, may suffer from aberrant immune responses^13^, ^14^, and can result in signaling discrepancies due to the lack of coreceptor stimulation^15^. To overcome these complications, the OT-1 TCR has been characterized and developed as a more physiological model system to study antigen-specific TCR signaling^16–18^. The OT-1 TCR recognizes the chicken ovalbumin-derived peptide OVA257-264 (SIINFEKL) bound to MHC tetramers as an agonist with strong binding affinity^19, 20^, facilitated by the interaction of coreceptor CD8 with MHC^21^. A panel of altered peptide ligands of OVA with sequentially reduced binding affinities have also been subsequently characterized^20, 22^. In our efforts to better understand T cell signaling, antigenic pMHC tetramer might be a more relevant activating agent compared to anti-TCR antibody. Physiologically, antigen affinities could lead to different biological outcomes for T cells because high-affinity pMHC induces negative selection while low-affinity pMHC leads to positive selection during thymic development^23^. Mechanistically, differences in ligand affinities also form the basis of the kinetic proofreading model in TCR initiation by manifesting in distinct binding kinetics^24^, ^25^. Due to the lack of wide-scale pTyr proteomic studies directly comparing pMHC and anti-TCR antibodies, it remains enigmatic whether the limitations in antibody-based approach had resulted in discrepancies in pTyr-mediated signaling pathways that could invalidate previous findings in T cell biology.

Furthermore, monoclonal anti-TCR antibodies^26^ and pMHC tetramers^27^ have become invaluable immunomodulating agents in conjunction with tyrosine kinase-targeting drugs^28^ for immunotherapy. However, methods to systematically evaluate phosphorylation-driven perturbations by these clinically relevant immunomodulators have not been fully established.

Here, we provide a framework based on the BOOST approach to systematically evaluate tyrosine phosphorylation-driven perturbations by anti-TCR antibodies and pMHC tetramers in an unbiased manner. We use the recently engineered Jurkat T cell line expressing OT1 TCR and CD8 coreceptor as an antigen-specific model TCR system to provide a readout of TCR signaling based on varying affinities of OVA pMHC altered peptide ligands. We also directly compare it with the pTyr-mediated signaling pathways triggered by antibody-mediated activation of TCR. For the first time, we demonstrate the utility of BOOST as a pTyr proteomics method to reveal biological insights mediated by tyrosine phosphorylation in T cells. Collectively, this approach allows the reproducible quantification of close to a thousand pTyr sites collectively in this dataset. We evaluate the perturbations in signaling pathways attributable to these pTyr sites and analyze the similarities and differences between antigen-specific activation of OT-1 receptor and antibody-directed stimulation. Importantly, we highlight how we optimize experimental design of BOOST and vigorously evaluate the data quality. Finally, we demonstrate the potential applications of BOOST in expanding our understanding of disease mechanisms and discuss advantages of the affinity-tunable antigen-specific pMHC to allow for a more precise control of TCR signaling strength.

## Experimental Methods

### Cell line and cell culture

Jurkat OT1+ CD8+ (J.OT1) cells were generated using lentiviral and retroviral vectors as described in detail previously^21^. Briefly, the human leukemic Jurkat T cell line (clone E6-1) that was made Lck-deficient using CRISPR-Cas9 was subsequently transduced with OT-I TCRβ and OT-I TCRα, hCD8β-T2A-hCD8α and Lck with a C-terminal FLAG sequence.

J.OT 1 cells were maintained in RPM1 1640 containing 2.05 mM L-glutamine supplemented with 100 U/mL penicillin G, 100 μg/mL streptomycin, 2 mM L-Glutamine (HyClone) and 10% (v/v) heat-inactivated fetal bovine serum (Peak Serum), in a humidified incubator with 5% CO_2_ at 37°C. Cells used in this work were confirmed mycoplasma-free using the Mycoplasma PCR Detection Kit (Applied Biological Materials G238) according to manufacturer’s instruction.

### T cell receptor stimulation

The stimulation method of J.OT1 cells is based on a previously established study^21^. For each replicate per condition, 15 million J.OT1 cells were washed and resuspended in serum-free RPMI at a concentration of 50 million cells per mL. Cells were then incubated with either 10 nM of H-2K^b^ MHC tetramer (Class I, human beta-2-microglobulin) or ~1 ug/mL of anti-CD3 IgM antibody C305 on ice for 1 h. C305 was diluted in Dulbecco’s phosphate-buffered saline (DPBS) prior to stimulation. MHC tetramers^29^ were synthesized by the NIH Tetramer Core Facility (Atlanta, GA) using custom peptides from GenScript. The peptide sequences are SIINFEKL (OVA), SIITFEKL (T4), SIIGFEKL (G4), and RGYVYQGL (VSV). Stimulation was initiated by transferring the cells from ice to 37°C. After 3 mins of stimulation at 37°C, cells were lysed with equal volume of lysis buffer (pH 7.6) containing 1% (w/v) sodium dodecyl sulfate (SDS), 100 mM Tris-HCl and 1X MS-SAFE Protease and Phosphatase Inhibitor cocktail (Sigma-Aldrich MSSAFE). PV treatment was performed by incubating J.OT1 cells with 500 μM PV (prepared by mixing equal volume of 1 mM sodium orthovanadate and 1 mM hydrogen peroxide) for 10 minutes at 37°C and lysed with the lysis buffer as mentioned.

### Sample processing for proteomic analysis

Lysate was applied through QIAshredder Mini Spin Column to reduce the viscosity of lysate by centrifugation at 20,000 x g at 37°C for 5 mins. Protein concentration of the clarified lysate was determined using Pierce BCA Protein Assay (Thermo Fisher Scientific, 23225), after which it was reduced by 100 mM dithiothreitol at room temperature for 30-60 mins. Lysate was subsequently processed using the filter-aided sample preparation (FASP) method^30^ as described. Briefly, 8 M urea in 100 mM Tris-HCl, pH 8.5 (UA) was used to dilute the SDS concentration of lysate to no more than 0.2 % (w/v). Lysate containing 200 ug of protein was applied to each Microcon-30 Ultracel PL-30 Regenerated Cellulose 30000 NMWL (Millipore, MRCF0R030) filter unit (1 mg of protein per replicate using 5 filter units in total) via centrifugation. 50 mM of iodoacetamide in UA was added to the filter unit and incubated in dark for 30 mins to alkylate the protein. Filter units were subsequently washed by UA and 50 mM ammonium bicarbonate, before the addition of trypsin (Promega, V5113) at 37°C overnight at a trypsin:protein ratio (w/w) of 1:40. Digested peptides were collected and acidified by trifluoroacetic acid (TFA) and desalted using Sep-Pak C18 Cartridge (Waters WAT020515) as described^31^.

### TMT labeling

The BOOST method was recently introduced^6^ and adopted in this study with slight modifications. Desalted peptides were labeled using a Tandem Mass Tag 11-plex isobaric label reagent set (ThermoFisher #A34808) with the following setup. Cells treated with PV were used as samples in the Boost channel (TMT126) across all TMT mixes. To prevent reporter ion interference from the ^15^N and ^13^C isotopes of TMT126 which can negatively impact quantitation^32^, TMT127N and TMT127C were left blank and unused. The rest of the sample channel setup are detailed in Figure 1C. To initiate TMT labeling, 1250 μg of TMT label (resuspended in 63 μL of acetonitrile) was added to peptides (resuspended in 150 μL of 100 mM triethylammonium bicarbonate) originating from 1 mg of protein for 3 h at room temperature. The labeling reaction was quenched by the addition of 12 μL of 5% (v/v) hydroxylamine for 15 mins at room temperature. To account for variation in peptide abundance, a portion of individually labeled peptides from each TMT channel within the same mix were pooled equally for a crude total peptide analysis. Briefly, a small aliquot (~2%) of individual TMT-labeled peptides was pooled and injected to the LC-MS without fractionation and searched using fixed modification of TMT11 while keeping all other parameters identical as described below. A separate search using variable modification of TMT yielded an average TMT labeling efficiency of ~99% across all four TMT mixes. Using data from the fixed modification search, median intensities of each of these TMT channels were used to normalize for pooling the remaining ~98% TMT-labeled peptides in equal amounts. The acetonitrile concentration of the pooled peptide mixture was reduced to less than 2% (v/v) and acidified to 1% TFA (v/v), prior to desalting using Sep-Pak C18 Cartridge (Waters WAT020515) and freeze-dried on a lyophilizer before proceeding with pTyr peptide enrichment as described^6^.

**Figure 1:**
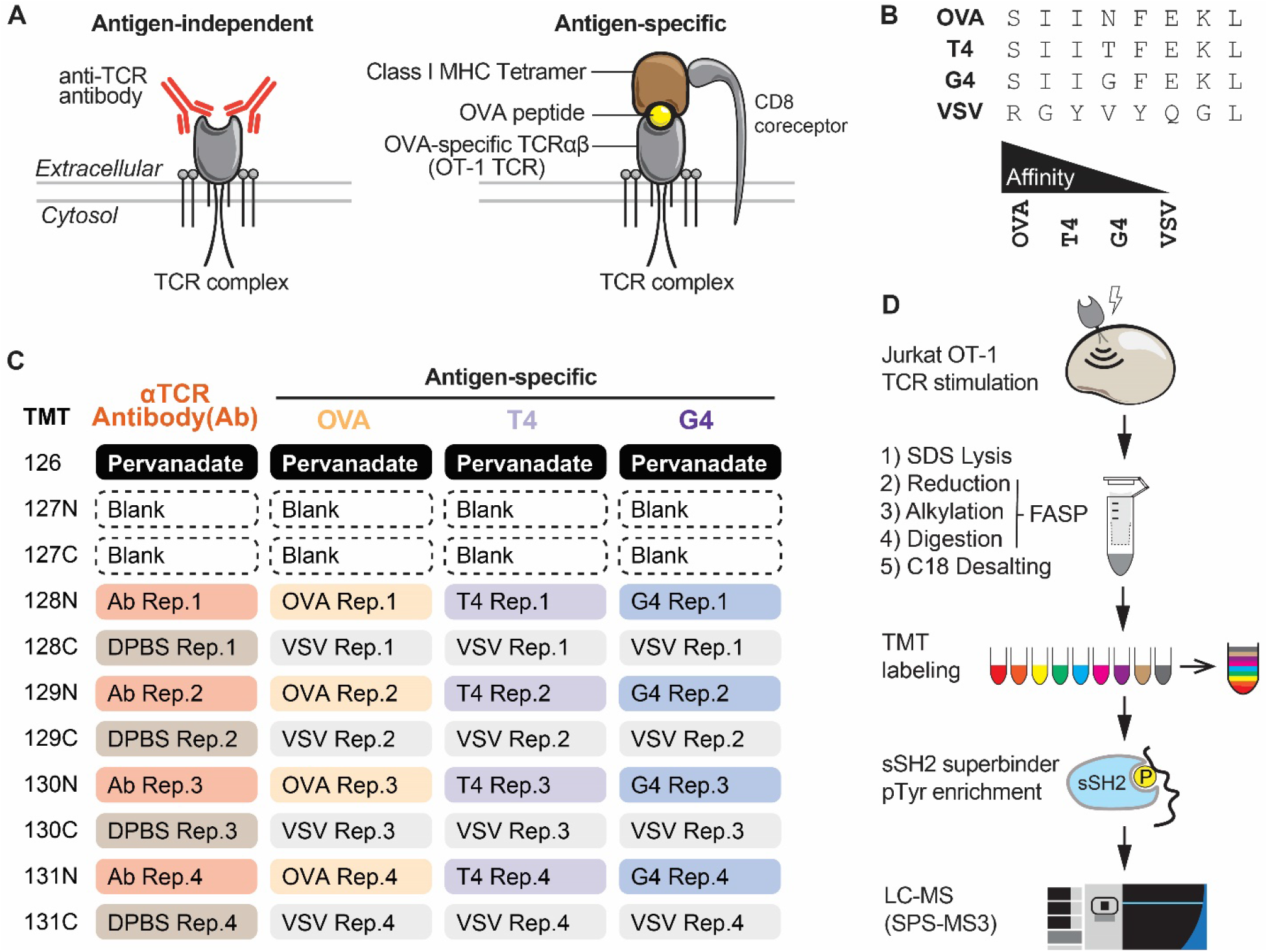
(A) Illustration of the differences between antigen-independent and antigen-specific triggering of TCR. Here, anti-TCR antibody is used for antigen-independent activation while the OVA pMHC tetramer targeting OT-1 TCR is used for antigen-specific activation of T cell signaling. For clarity, MHC is depicted as a monomer instead of a tetramer. (B) Amino acid sequence of various altered peptide ligands of the OVA peptide. Alteration of the peptide sequence reduces the binding affinity of the pMHC to OT-1 TCR. (C) Sample designation of TMT channels used in this study. A total of four TMT11 mixes were used. PV-treated samples were labeled with TMT126 as BOOST channel to increase the quantitation depth of the tyrosine phosphoproteome. For the comparison of Ab, DPBS was used as the unstimulated control because anti-TCR antibody was diluted in DPBS, while for the comparisons of OVA, T4 and G4 pMHC, null peptide VSV pMHC was used as their respective unstimulated control. (D) Overview of the experimental flow. Jurkat T cells expressing OT-1 TCR were stimulated as shown in (C) and lysed with SDS lysis buffer. Lysate was reduced, alkylated and digested using the filter-aided sample preparation (FASP) methods. Digested peptides were desalted on a reverse-phase C18 column and subsequently TMT-labeled prior to pTyr peptide enrichment using sSH2 superbinder. Proteomic data were collected on a mass spectrometer using the synchronous precursor selection (SPS) MS3 parameters. More details can be found on the Experimental Methods Section.

### pTyr peptide enrichment using SH2 superbinder

The Src SH2 domain superbinder was purified and immobilized on CNBr-activated Sepharose (GE Healthcare) in IAP buffer (50 mM MOPS-NaOH [pH7.2], 10 mM Sodium Phosphate, 50 mM Sodium Chloride) as described^4^ at a concentration of 2.4 mg protein per mL of beads slurry (50 mg Sepharose beads per mL). 9 mg of desalted TMT-labeled peptides were resuspended in 2.8 ml of IAP buffer and subsequently added to 2 mg of immobilized SH2 superbinder to be incubated on a rotator overnight at 4°C. The beads were washed 3 times with ice-cold IAP buffer and once with ice-cold HPLC-grade water. All washes and removal of bead supernatant during the enrichment procedure were performed using a centrifugation speed of 1,500 x g at 4°C for 2 mins unless stated otherwise. Peptides were eluted from the beads twice, each using 200 μL of 0.15 % (v/v) TFA for 10 minutes at room temperature with constant agitation. Eluted peptides were desalted using 100 μL C18 tips (Thermo Scientific Pierce) following the manufacturer’s guidelines and dried on a speed-vac.

### Liquid chromatography-mass spectrometry

For offline basic (pH 10) fractionation, peptides were separated on a 100 mm x 1.0 mm Acquity BEH C18 column (Waters) using an UltiMate 3000 UHPLC system (ThermoFisher Scientific) with a 40-minute gradient from 1% to 40% Buffer B_basic_ into 36 fractions, which are subsequently consolidated into 12 super-fractions (Buffer A_basic_ = 10 mM ammonium hydroxide in 99.5% (v/v) HPLC-grade water, 0.5% (v/v) HPLC-grade acetonitrile; Buffer B_basic_ = 10 mM ammonium hydroxide in 100% HPLC-grade acetonitrile). Each super-fraction was further separated on an in-line 150 mm x 75 μm reversed phase analytical column packed in-house with XSelect CSH C18 2.5 μm resin (Waters) using an UltiMate 3000 RSLCnano system (ThermoFisher Scientific), at a flow rate of 300 nL/min. Peptides were eluted using a 65-minute gradient from 5% to 30% Buffer B_acidic_, followed by a 6-minute gradient 30% to 90% Buffer Bacidic (Buffer Aacidic = 0.1% (v/v) formic acid in 99.4% (v/v) HPLC-grade water, 0.5% (v/v) HPLC-grade acetonitrile; Buffer B_basic_ = 0.1% (v/v) formic acid in 99.9% (v/v) HPLC-grade acetonitrile). Data was acquired in DDA mode on a Orbitrap Eclipse Tribrid mass spectrometer (ThermoFisher Scientific) with a positive spray voltage of 2.25 kV using multinotch TMT-MS3 settings^33^. Cycle time was set at 2.5 seconds. At the MS1 level scans, precursor ions (charge states 2-5) acquired on the Orbitrap detector with the scan range of 400-1600 m/z, 120,000 resolution, maximum injection time of 50 ms, automatic gain control (AGC) target of 800,000, and a dynamic exclusion time of 15 seconds. MS1 precursor ions were isolated on the quadrupole using an isolation window of 0.7 m/z for MS2 scans. MS2 scans were acquired in centroid mode on the ion trap detector on a scan range of 400-1400 m/z via higher-energy dissociation (HCD, 33% energy) activation with an AGC target of 5000, maximum injection time of 75 ms. Using synchronous precursor selection (SPS)^33^, 10 notches were further isolated on the quadrupole using an MS2 isolation window of 3 m/z for MS3 scans, which are acquired on the Orbitrap detector on a scan range of 100-500 m/z in a mass resolution of 50,000 via HCD activation (55% energy) with a AGC target of 250,000 and maximum injection time of 150 ms in centroid mode.

### Database Search Parameters and Acceptance Criteria for Identifications

Raw files were processed in MaxQuant^34^ version 1.6.17.0 using the integrated peptide search engine Andromeda^35^. MS/MS spectra were searched against a human UniProt database (*Homo sapiens*, last modified 12/01/2019) comprised of 74,811 forward protein sequences. False discovery rate (FDR) for peptide spectrum matches (PSM) was set at 1% using a reverse decoy database approach. Carbamidomethylation (cysteine) was set as fixed modification, whereas oxidation (methionine), acetylation (protein N-termini) and phosphorylation (serine, threonine, tyrosine) were set as variable modifications. Trypsin enzyme specificity was used with up to 2 missed cleavages. Main search peptide tolerance was set as 3ppm, while FTMS and ITMS MS/MS match tolerances were set as 20 ppm and 0.5 Da, respectively. MS3 reporter ion mass tolerance was set at 3 mDa, using isotopic correction factors provided by the manufacturer (Lot UK291565, Lot UH283151). Match between runs was enabled with the default parameters. As published previously^6^, imputation or interpolation of missing values was not used in the analysis of TMT data. Search parameter file (mqpar.xml) of the MaxQuant run is provided as File S1.

### Immunoblotting

To prepare cell lysate for immunoblots, cell pellets were resuspended in sample loading buffer (2% w/v SDS, 62.5 mM Tris-HCl, 10% v/v glycerol, 2.5% v/v 2-mercaptoethanol, 0.05% w/v bromophenol blue) and boiled at 95°C for 10 minutes. Approximately 500,000 cell equivalents or 25 μg of protein was loaded into each well of ClearPAGE 4-20% polyacrylamide gradient gels (Expedeon) and resolved by denaturing polyacrylamide gel electrophoresis at 100 Volts for 1 h. Proteins resolved by gel electrophoresis were transferred onto an Immobilon-FL PVDF membrane (EMD Millipore) at 100 Volts for 90 minutes in ice-cold transfer buffer (25 mM Tris, 200 mM glycine, 20% v/v methanol). Membranes were then blocked with Odyssey Blocking Buffer (Li-Cor) at room temperature for 1 h before incubation with primary antibodies at 4°C overnight and subsequent incubation with IRDye-conjugated secondary antibodies (Li-Cor) for 1 h at room temperature. Membranes were washed with DPBS containing 0.1% v/v Tween after each antibody incubation. Anti-GAPDH was from Sigma-Aldrich (G9545, 1:10000 dilution). Anti phosphotyrosine antibody (4G10, 05-321) was from Millipore. Anti-p44/42 MAPK (Erk1/2, 9107), anti-phospho-p44/42 MAPK (Erk1/2, Thr202/Tyr204, 9101), anti-Zap70 (3165), anti-Zap70 pTyr493 (2704) and anti-LAT pTyr191 (pTyr220 in isoform 1, 3548) antibodies were from Cell Signaling Technologies. IRDye 680RD-conjugated anti-rabbit IgG (926-68071) and IRDye 800CW-conjugated anti-mouse IgG (926-32212) secondary antibodies were from Li-Cor. All primary antibodies were diluted 1:1000 (v/v) while all secondary antibodies were diluted in 1:10000 (v/v) unless stated otherwise. Blots were visualized using the Odyssey CLx Imaging System and quantified using ImageStudio software (Li-Cor).

### Data Analysis

For PSM level analysis, “evidence.txt” (Table S1) generated from MaxQuant was used. Unique PSM is defined by a peptide spectrum match with non-redundant amino acid sequence, modifications and charge state with the least number of missing values across all TMT channels. If redundancy still exists, PSM with the highest median reporter ion intensity is retained. For pTyr level analysis, “Phospho (STY)Sites.txt” (Table S2) from MaxQuant was used. After removing reverse hits and potential contaminants, only Class I pTyr sites (localization probability >0.75) were included in the analysis. Reporter ions from TMT126 (PV-treated samples) and TMT127N-TMT127C (blanks) were excluded from all quantitative analyses, unless stated otherwise. Unpaired student’s t-test was performed using the corrected phosphosite intensities between up to 4 replicates of each paired condition within the same TMT mix to determine the p values, of which are adjusted to q values to account for FDR in multiple testing hypothesis^36^. Volcano plots were constructed using -log_10_(q values) and log_2_(fold change in mean pTyr site intensities) of the stimulated condition versus its corresponding control as detailed in Figure 1C, with a requirement that at least 3 out of 4 replicates for each of the pairwise condition for the same pTyr site contained a reporter ion intensity greater than zero. As published previously^6^, imputation or interpolation of missing values was not used in any part of the data analysis. pTyr sites with q values less than 0.05 were considered statistically significant. The pTyr sites were also annotated to a KEGG database based on the Uniprot accession number of the corresponding protein using Perseus^37^. Intensity heatmap of pTyr site fold change was generated after performing hierarchical clustering based on the Euclidean distances between cluster means using the complete linkage method. Due to the length of the ratio heatmap of the entire dataset, an interactive heatmap is provided and is available to be examined interactively online (details on File S2). For PTM signature enrichment analysis (PTM-SEA), we adopted the methodology developed by Krug et al.^38^ and R scripts from Storey et al.^39^ using 7 amino acids flanking each pTyr site with the following modifications. The input for PTM-SEA was modified to be a vector of log-transformed q values multiplied by the sign of the mean log2 reporter ion ratio, instead of p values. The resulting formula is *-log10(q value) * sign [log2(fold change in pTyr site)]*. To fully capture the variance observed across replicates, we deployed a similar approach as Krug et al. to incorporate the magnitude of phosphosite q values into the determination of enrichment scores for each of the signature categories, with no rank normalization and by setting the weight parameter to 1^38^. Venn diagram was generated using InteractiVenn. All analyses and plotting were performed in Microsoft Excel or using the statistical language R (4.0.3) in RStudio (1.1.447) using packages “tidyverse” (1.3.0), “qvalue” (2.22.0), “ggdendro” (0.1.22), “plotly” (4.9.3), “RColorBrewer” (1.1-2), ““pacman” (0.5.1), “rhdf5” (2.34.0), and “cmapR” (1.2.1) downloaded from CRAN or Bioconductor. All R scripts used in this study are available to be downloaded in (Files S3-6). Immunoblot quantitation is provided in File S7. KEGG-annotated pTyr sites can be found in Table S3.

## Results

### BOOST Experimental Design and Rationale

We set out to investigate the pTyr signaling pathways triggered by antigen-independent or antigen-specific activation of TCR by deploying the BOOST approach we recently developed. Briefly, PV-treated sample(s) in a TMT-multiplexed BOOST strategy enables deeper pTyr quantitative depth by selectively triggering the fragmentation of pTyr-containing precursor ions, thereby facilitating reporter ion quantitation of samples-of-interest. We verified efficacy of PV by examining the elevation of total tyrosine phosphorylation levels across the proteome of PV-treated samples in an immunoblot (Figure S1). We used Jurkat T cells expressing OT-1 TCR and CD8 coreceptor as a model system for antigen-specific activation of TCR (Figure 1A) because the OT-1 TCR has been demonstrated to bind H-2K^b^ OVA pMHC class I tetramers with high affinity^19^, ^20^. Additionally, a panel of OVA peptide variants with sequentially reduced binding affinity has been characterized, of which we investigated the T4 (medium affinity), G4 (low affinity) and VSV (null peptide control) antigens in this study (Figure 1B). To minimize discrepancies in signaling due to heterogeneity arising from different cell lines, we used the same Jurkat OT-1 cell line for all conditions in this study because this cell line can also be stimulated by C305 IgM antibody targeting TCR Vbeta^10^, ^21^ in an antigen-independent manner (Figure 1A). This allows for a more direct comparison of TCR signaling readout solely from the activating agent.

To quantitatively characterize the differences in signaling pathways induced by these activating agents, we performed a multi-batch TMT BOOST experiment to probe the changes in pTyr sites as a result of TCR activation by anti-TCR antibody (Ab), OVA pMHC (OVA), T4 pMHC (T4) and G4 pMHC (G4) in quadruplicates each (Figure 1C). Appropriate controls were chosen for each of these conditions. Because Ab was diluted in DPBS in this experiment, DPBS was used as the no-Ab control, while the unrelated vesicular stomatitis virus pMHC (VSV) was used as the null-peptide controls for OVA, T4 and G4 pMHCs. TMT126 was designated as PV boost channel followed by 2 blank channels in TMT127N and TMT127C to avoid reporter ion interference of isotopic impurities from the TMT labeling reagent^32^ (Figure 1C). Samples were TMT-labeled and enriched for pTyr-containing peptides using sSH2 superbinder prior to MS3-based TMT reporter quantitation (Figure 1D). The selectivity of pTyr enrichment was demonstrated as the majority of phosphorylation sites identified was localized on tyrosine residues, with 94% of the pTyr sites categorized as Class I sites by MaxQuant (Figure S2).

### Data Quality and Precision were Evaluated

To ensure that the biological interpretation of the data is not negatively impacted by PV boost channel, we first inspected several metrics to assess the overall quality of the data. As expected from a BOOST experiment, the inclusion of PV boost channel resulted in a higher number of quantifiable unique PSM with ~7-10X reporter ion intensities in TMT126 compared to other TMT channels; this effect is specific to only PSMs with pTyr modifications across all conditions (Figure S3A-D). The strategic placement of 2 blank channels allowed us to circumvent the reporter ion interference due to isotopic impurities from PV boost channel (Figure S3A-D). Importantly, the inclusion of PV boost channel still resulted in good quantitative precision, with median coefficients of variation (CV) of less than 20% across all conditions for pTyr-containing PSMs of the biological replicates performed in this study (Figure S4). We attribute the observed quantitative precision to median PV reporter ion boost levels of a modest 4-to 30-fold relative to the average experimental reporter ion intensity of pTyr-containing PSM, depending on the number of missing values (Figure S5). Importantly, boost levels for PSM without pTyr modification(s) remain unchanged throughout, suggesting that the effect of PV boost channel is highly specific and targeted (Figure S5). A more detailed interpretation of these data is provided in the Discussion section. Altogether, the overall data quality and good quantitative precision provided confidence in the interpretation of our subsequent data analysis.

### OVA Triggered a Similar but Potentially Stronger Signaling Relative to Ab

To examine the changes in abundance of pTyr sites in relation to statistical significance, we visualized the fold changes of each quantified pTyr site in volcano plots (Figure 2A-E). To maintain statistical stringency, we required a minimum of 3 out of 4 replicates with quantifiable reporter ions for each condition within the comparison to allow for the calculation of q value (FDR-corrected p value), resulting in a total of 996 pTyr sites collectively that can be examined using volcano plot analysis (Figure 2F). pTyr sites with a q value less than 0.05 are considered statistically significant or differential. Among these differential pTyr sites, many of these sites are found in proteins known to be activated and phosphorylated upon T cell activation, notably CD3 and CD247(ζ) chains forming part of the TCR complex. Not only did we identify all ITAM pTyr sites within CD3 and CD247(ζ) chains (CD247 Y72/Y83, Y111/Y123, Y142/Y153; CD3D Y149/Y160; CD3E Y188/Y199; CD3G Y160/171), our data also revealed that nearly all of these ITAM pTyr sites were statistically increased in both Ab and OVA upon TCR activation (Figure 2A-C, Figure 3A). These similarities in differential ITAM sites suggest that a comparable early signaling machinery between Ab and OVA, although the magnitude of ITAM phosphorylation were mostly larger for strong agonist OVA (Figure 3A). For instance, OVA triggered up to 6-fold increase in ITAM sites but only up to about 3-fold increase was observed for Ab (Figure 3A). Moreover, the majority of the fold changes in pTyr sites involved in TCR signaling that were induced by OVA were also higher than that of Ab (Figure 3B), indicative of a stronger signaling output overall. It is possible that a stronger effect was observed in OVA-activated samples due to the contributions of CD8 coreceptor which binds specifically to MHC. These data are consistent with the overlap in differential sites between Ab and OVA, where 27 out of 40 (or approximately 7 in 10) pTyr sites that were identified as statistically significant in Ab were also similarly identified in OVA, but OVA had ~1.7X more differential pTyr sites than Ab (Figure 2F). Interestingly, OVA-specific differential pTyr sites also included several well-known negative TCR regulators such as UBASH3A, PAG1 and PTPN6 (SHP-1) (Figure 2A, C), possibly indicating the initiation of a negative feedback inhibitory regulation of strong signaling induced by OVA. Taken together, our data suggest a close resemblance in early pTyr signaling between antigen-independent Ab and antigen-specific OVA pMHC activation of TCR, but concurrently indicate a stronger signaling output induced by the high-affinity antigen OVA.

**Figure 2:**
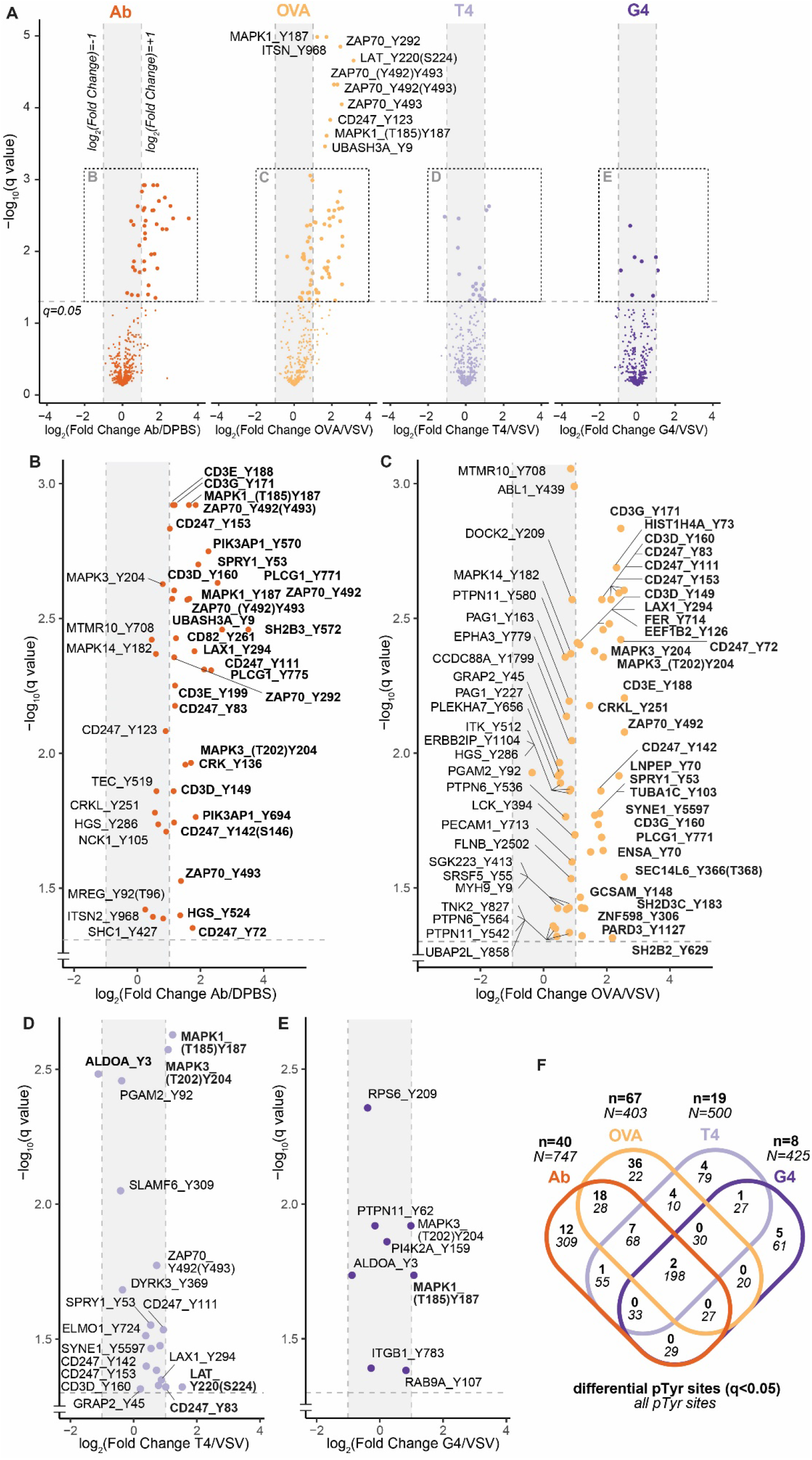
(A) Volcano plots quantifying the fold changes of pTyr sites in all four TMT comparisons as shown in Figure 1C, for pTyr sites with at least 3 out of 4 channels with quantified reporter ions in each condition of the comparison. Vertical dashed lines represent 2-fold increase or decrease in the abundance of pTyr site. Horizontal dashed line depicts a q value cutoff of 0.05. pTyr sites with q values lower than 0.05 are considered statistically significant. Significant pTyr sites are labeled using the notation [gene_name]_[Y][residue_number] (gene name is used instead of protein name for brevity). pTyr sites originating from doubly phosphorylated peptides are annotated with a parenthesis containing the second phosphorylation site with the highest localization probability. pTyr sites without a parenthesis originate from singly phosphorylated peptides. To allow better visibility, significant pTyr sites in the boxed regions are scaled and labeled separately in (B-E). (B-E) Significant pTyr sites (q<0.05) are shown and labeled for comparisons of Ab (B), OVA (C), T4 (D) and G4 (E). To further bin significant pTyr sites, sites that are greater than or less than 2-fold change are highlighted in bold. (F) Venn diagram showing the number of overlapping pTyr sites identified in the volcano plots in (A) between each condition. The number of differential (q<0.05) pTyr sites (**n**) is shown in bold at the top while the number of all pTyr sites (*N*) is shown in italic at the bottom. A minimum of 3 out of 4 replicates with quantifiable reporter ions for each condition within the comparison was required to be reported in *N*.

**Figure 3:**
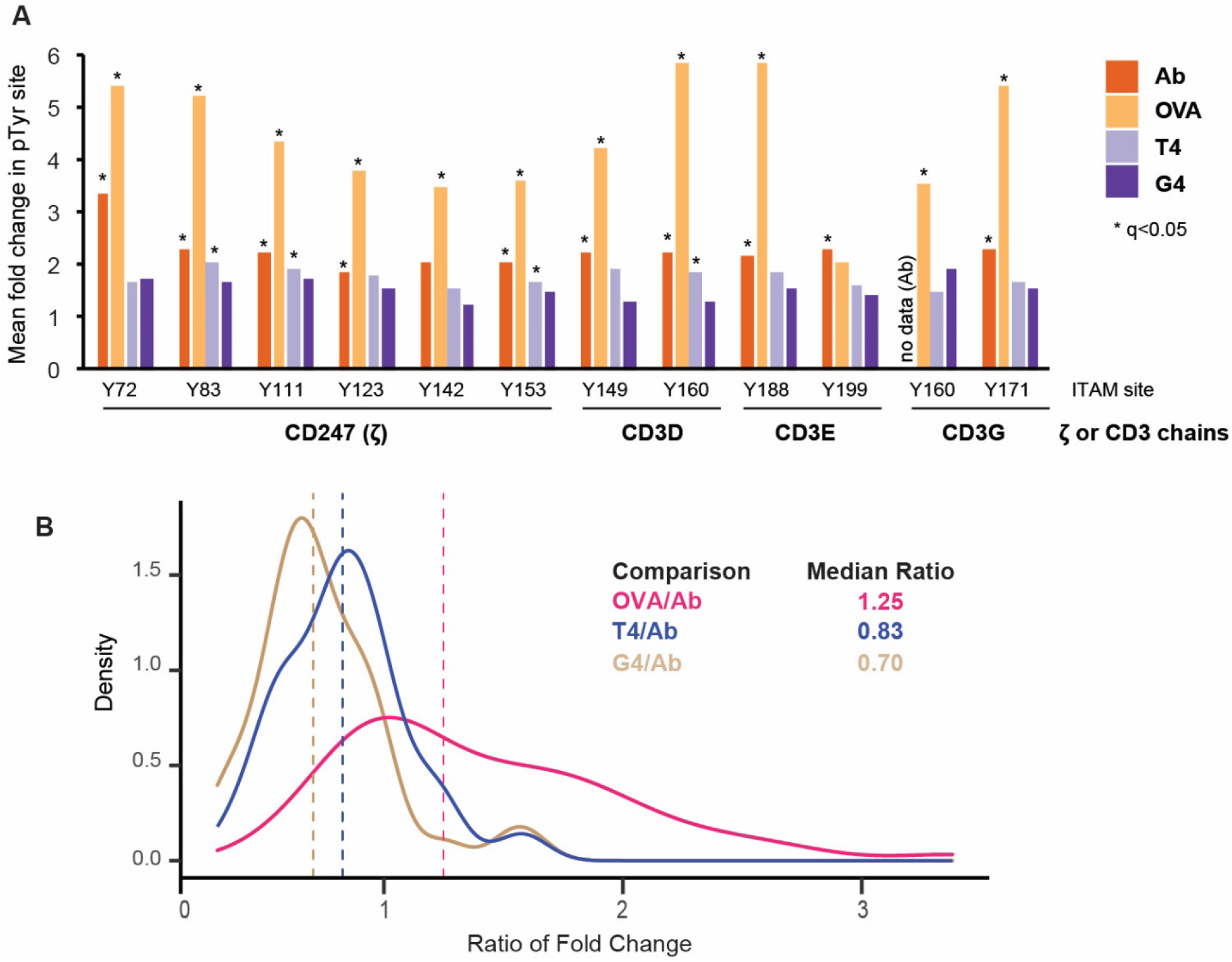
(A) The mean fold changes in pTyr intensities of ITAM sites in ζ and CD3 chains are shown for each stimulated condition relative to its corresponding unstimulated control as detailed in Figure 1C. * above the bar indicate statistically significant site defined by a q value of less than 0.05. (B) Density plot showing the distribution of the ratio of fold changes in paired pTyr site intensities (KEGG TCR signaling proteins) in OVA/VSV, T4/VSV and G4/VSV relative to Ab/DPBS. Dashed vertical line indicate the median of all ratios of fold changes, which is also numerically indicated on the plot. Here, a ratio of fold change of 1 indicates that the fold change in a pTyr site induced by a pMHC is equivalent to that of the fold change in the same pTyr site induced by Ab, relative to their corresponding unstimulated control, respectively.

### Signaling Strength Induced by PMHC was Consistent with Binding Affinity

To examine if the predicted binding affinity of OVA, T4 and G4 pMHC (Figure 1B) can be reflected in a readout of TCR signaling strength induced by tyrosine phosphorylation, we next compared the differential pTyr sites elicited by these antigens. Indeed, the overall magnitude in T cell signaling strength driven by the changes in tyrosine phosphorylation was reflected by the fold changes in ITAM sites and the relative fold changes in pTyr sites involved in TCR signaling (Figure 3A-B), which is consistent with the predicted binding affinities of the pMHC. While all but one ITAM pTyr sites were statistically significant in OVA, only 4 were differential in T4 and none in G4 (Figure 3A). Furthermore, a total of 67 pTyr sites were statistically significant in OVA, while only 19 and 8 differential pTyr sites were identified in T4 and G4 respectively, despite comparably similar number of total pTyr sites analyzed in each condition (Figure 2F). For instance, the activating pTyr sites of many characterized early TCR signaling proteins, such as Lck Y394^40^, Zap70 Y493^41^, LAT Y220^42^, ITK Y512^43^, and PLCG1 Y771^44^, were statistically increased in the high-affinity antigen OVA, but not in the medium- or low-affinity antigens T4 and G4 (Figure 2A, 2C-E). However, these antigens generally triggered a similar but weaker signaling output, since we still observed a small but noticeable induction of ZAP70, LAT, CD3 chains and CD247 pTyr sites in T4 (Figure 2D). Collectively, the discrepancy in differential pTyr sites between OVA, T4 and G4 showed how effectively BOOST can discriminate between pMHCs of varying binding affinities based on the readout of TCR signaling strength induced by tyrosine phosphorylation.

### Validation of Proteomic Data Revealed Similarities in Downstream MAPK Activation Pathways

To demonstrate that the quantitation from the BOOST proteomic data accurately reflects the cellular state of our experimental samples, we sought to validate our proteomic data using an alternative approach, such as immunoblot. Interestingly, only 2 pTyr sites were statistically increased in all 4 conditions, which are MAPK1 (Erk2) (T185)Y187 and MAPK3 (Erk1) (T202)Y204 (Figure 2A-F). Activation of the canonical TCR signaling is known to activate the mitogen-activated protein kinase (MAPK) cascade, which sequentially activates Ras, Raf and MEK^45^. MEK activate MAPK1 and MAPK3 by dual phosphorylation of a conserved tripeptide TxY motif^46^. We used a phosphosite-specific antibody that recognizes the dual phosphorylation sites of MAPK1 and MAPK3 using immunoblot to validate the quantitation from our proteomic data (Figure 4A). Encouragingly, quantitating the fold changes of these dual phosphosites by immunoblot or by proteomic analysis were in full agreement with each other (Figure 4B). The data revealed largely comparable fold changes in MAPK1 and MAPK3 activation sites between Ab and OVA, likely suggesting that a similar MAPK-mediated pathway is activated downstream of the early ITAM signaling between Ab and OVA. Congruent with the results in ITAM sites (Figure 3A), average fold changes in MAPK1 and MAPK3 phosphosites for T4 and G4 correlated with their lower binding affinity relative to OVA (Figure 4B), consistent with a previous report^23^. Altogether, we validated the quantitation of our proteomic data in MAPK1 and MAPK3 phosphorylation sites and revealed potential similarities in downstream MAPK signaling pathways between antigen-independent Ab and antigen-specific OVA pMHC.

**Figure 4:**
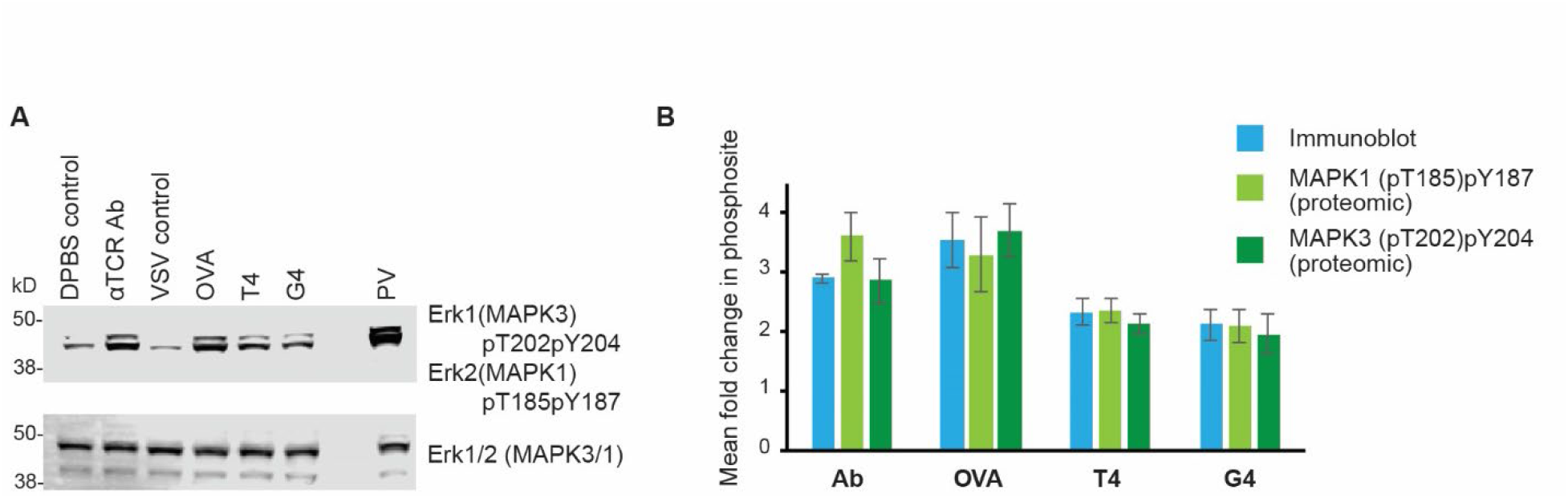
(A) Immunoblot of Erk1 (MAPK3) pThr202/pTyr204 and Erk2 (MAPK1) pThr185/pTyr187 phosphosites and total Erk1 and Erk3 proteins acting as loading controls. A representative image from 3 separate Western blots is shown. (B) Quantitation of MAPK1 and MAPK3 phosphosite fold changes of immunoblot compared to quantitation from proteomic data. DPBS was used as the unstimulated control for antibody (Ab), while VSV was used as the unstimulated control for OVA, T4 and G4. For immunoblot quantitation, band intensity for the dual phosphosites for both MAPK1 and MAPK3 was combined, averaged, and controlled by the loading controls, while the dual phosphosites for MAPK1 and MAPK3 were quantified separately for the proteomic data as indicated. Western blot was performed in triplicates; proteomic data was collected in quadruplicates. Error bar represents the standard deviation of the mean fold change.

### Phosphosite-centric Signature Enrichment Analysis using PTM-SEA

Instead of inspecting each pTyr site individually, a global phosphosite analysis of pTyr sites across all conditions would enable a more insightful interpretation of the cellular signaling triggered by various activating agents. To achieve this goal, we performed a global pathway analysis using PTM Signature Enrichment Analysis (PTM-SEA) built on a curation of modification site-specific databases such as PhosphoSite Plus and NetPath^38^. PTM-SEA allows for pathway analyses at the phospho-site centric level instead of at the gene-centric level to better incorporate the information from phosphosites instead of the gene products alone, which is ideal for a pTyr proteomic dataset. Using the PTM-SEA methodology to investigate perturbation signatures, we detected highly enriched signatures of anti-CD3 pathway across all conditions (Figure 5A). A positive enrichment score indicates a correlation between the annotated site and the signature category, while a negative score indicates an anti-correlation. Interestingly, anti-CD3 pathway is known to be triggered by many T cell activating antibodies^47^, ^48^, and this perturbation is similarly enriched by antigen-specific OVA, T4 and G4. Other perturbations enriched by OVA and T4 such as vanadate^49^, thrombin^50^, phorbol ester^51^, lipopolysaccharide (LPS)^52^, insulin^53^, IL-2^54^ were previously validated by antibody-based studies. Notably, the only perturbation with a negative enrichment score cross all conditions is dasatinib (Figure 5A), presumably because dasatinib is a selective tyrosine kinase receptor inhibitor used to treat chromic myeloid leukemia^55^, therefore is expected to have an anti-correlation with increased tyrosine phosphorylation upon TCR triggering. PTM-SEA also revealed similar sets of substrates for the kinases enriched in all conditions, notably Zap-70 and Lck (Figure 5B). Tyrosine kinases Lck and Zap-70 are known to be among the first kinases to be activated and recruited to the TCR complex upon T cell stimulation^56^. Known substrates of Lck or Zap-70 include ITAMs of CD3 and ζ (CD247) chains, LAT, and ITK, which were identified with increased abundance in pTyr sites in this dataset (Figure 2A-E). Interestingly, analysis using the NetPath database reported an enrichment of the prolactin pathway, with a trend of enrichment scores seemingly correlating with the strength of the antigen affinities of OVA, T4 and G4 (Figure 5C). The activation of human prolactin receptor by the prolactin hormone has been discovered to share many phosphorylation-driven signaling components with activated TCR pathways^57^, ^58^. Taken together, widescale examination of all pTyr sites using PTM-SEA revealed largely similar enrichment of signatures between antigen-independent and antigen-specific activation of T cell signaling.

**Figure 5:**
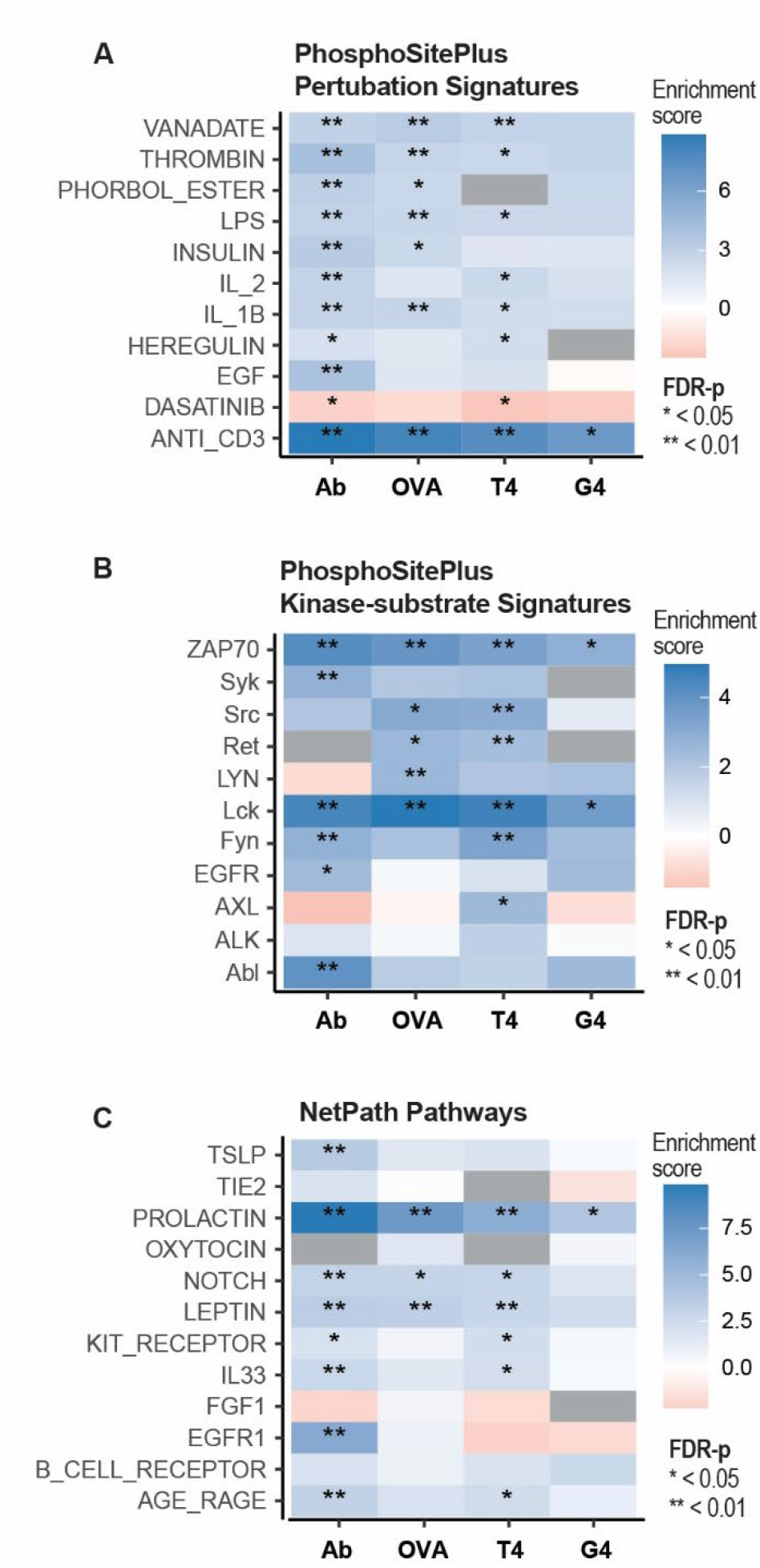
PTM signature enrichment analysis (PTM-SEA) of all pTyr sites with at least 3 out of 4 channels with quantifiable reporter ions in each condition of the comparison. The signed q values of each pTyr site were ranked ordered and used to calculate the enrichment score of each signature category known to correlate with the increase or decrease in abundance of the pTyr site based on PTMsigDB. PTMsigDB contains annotation of each pTyr site with the direction of regulation in various signature sets mined from PhosphoSitePlus perturbation signatures (A), PhoshosSitePlus kinase-substrate signatures (B), and NetPath Pathways (C). A positive enrichment score indicates a correlation between the annotated site and the signature category, while a negative score indicates an anti-correlation. Grey box means an enrichment score could not be calculated for that signature set in the comparison.

### Differential pTyr sites in Ab and OVA were Similarly Represented in Top 25 KEGG Pathways

To further assess whether the pathways reflected by differentially changing pTyr sites are consistent with the results from PTM-SEA above, we performed a non-parametric analysis using an alternative pathway database KEGG using only pTyr sites that were statistically significant (q<0.05) instead of all pTyr sites. The annotated KEGG pathways of the proteins containing the differential pTyr sites were parsed and ranked-ordered to obtain the top 25 KEGG pathways most represented by the differential pTyr sites (Figure 6). Predictably, the majority of these KEGG pathways were involved in the immune system, such as TCR signaling pathway, natural killer cell cytotoxicity, Chagas disease, and primary immunodeficiency. We observed that the number of differential pTyr sites were proportionally similar between Ab and OVA in almost every KEGG pathway analyzed, while the data for OVA, T4 and G4 seemingly correlate with the predicted affinity strength of respective pMHC. Collectively, the data suggest a close resemblance in signaling pathways implicated by antigen-independent Ab stimulation and high-affinity OVA antigen-specific activation of the TCR, while also demonstrate the sensitivity to discriminate between pMHC of weaker affinities (T4, G4) and their signaling output.

**Figure 6:**
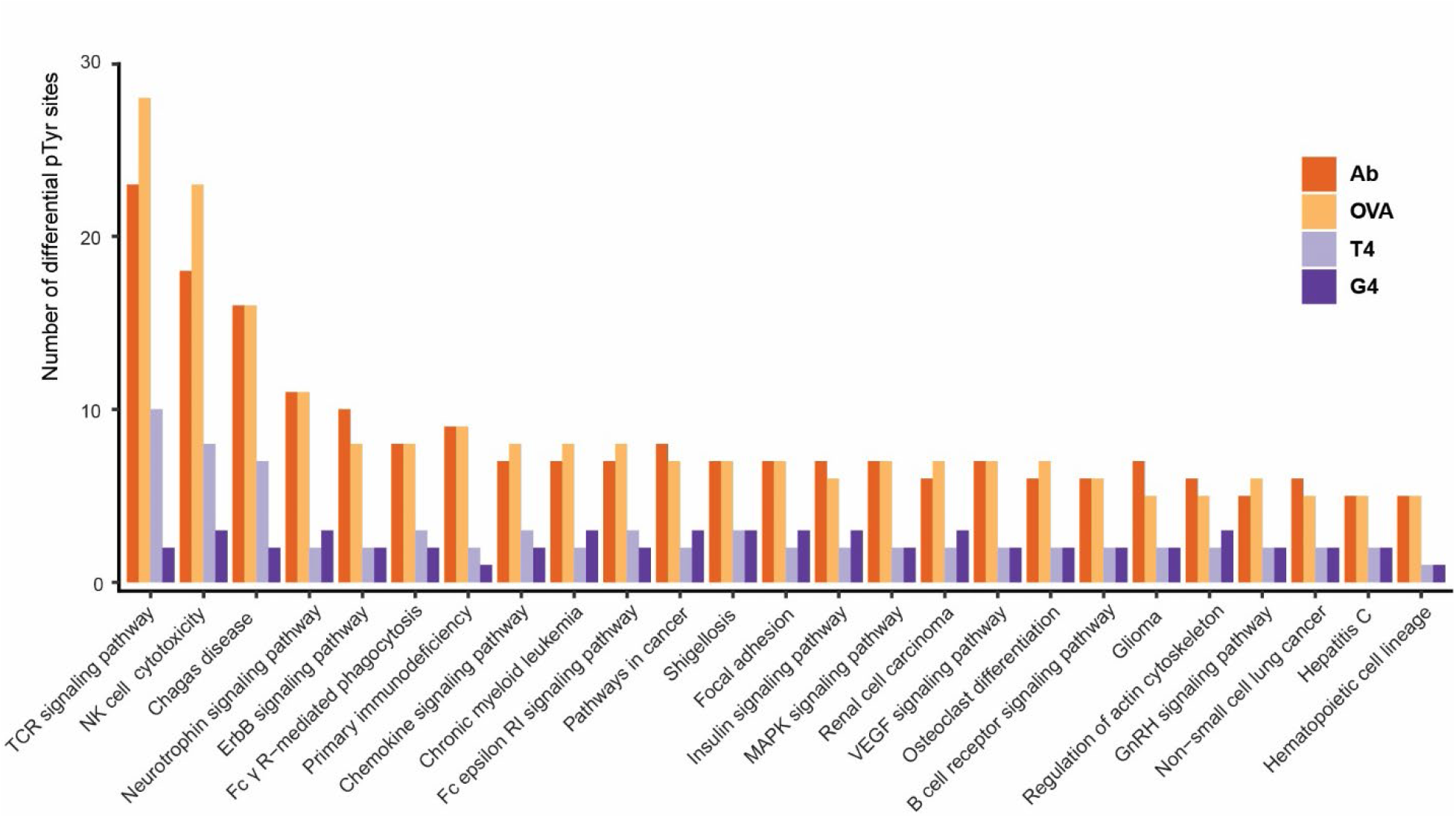
The number of significantly changing pTyr sites (q<0.05) in the top 25 KEGG pathways most represented by these differential pTyr sites across all comparison in this study.

### KEGG TCR Signaling Pathway Proteins Suggest Possible Negative Regulation in OVA

To further examine the proteins known to be involved in the T cell biology, we inspected all pTyr sites of the proteins that are involved in the TCR signaling pathway as annotated by KEGG. We compared the fold-change intensity in the form of a heatmap after performing hierarchical clustering (Figure 7). Similar clusters of elevated fold changes in pTyr sites were observed across all conditions, especially in canonical TCR signaling proteins such as CD3D, CD3E, CD3G, CD247, ZAP70, LAT, MAPK1 and MAPK3. Not only were the global fold-changes for the high-affinity antigen OVA the highest among all conditions, the data also included statistically increased Y536 and Y564 phosphorylation sites of PTPN6 (SHP-1) which were not observed in other conditions. This is notable because PTPN6 is a tyrosine phosphatase that negatively regulates T cell signaling, and the phosphorylation of Y536 and Y564 were reported to correlated with increase phosphatase activity^59^. Overall, the data suggest largely similar profile in the pTyr sites of proteins involved the KEGG T cell signaling pathway between antigen-independent Ab stimulation and antigen-specific activation of the TCR, but high-affinity OVA showed possible signs of a negative feedback regulation likely due to a stronger signaling effect.

**Figure 7:**
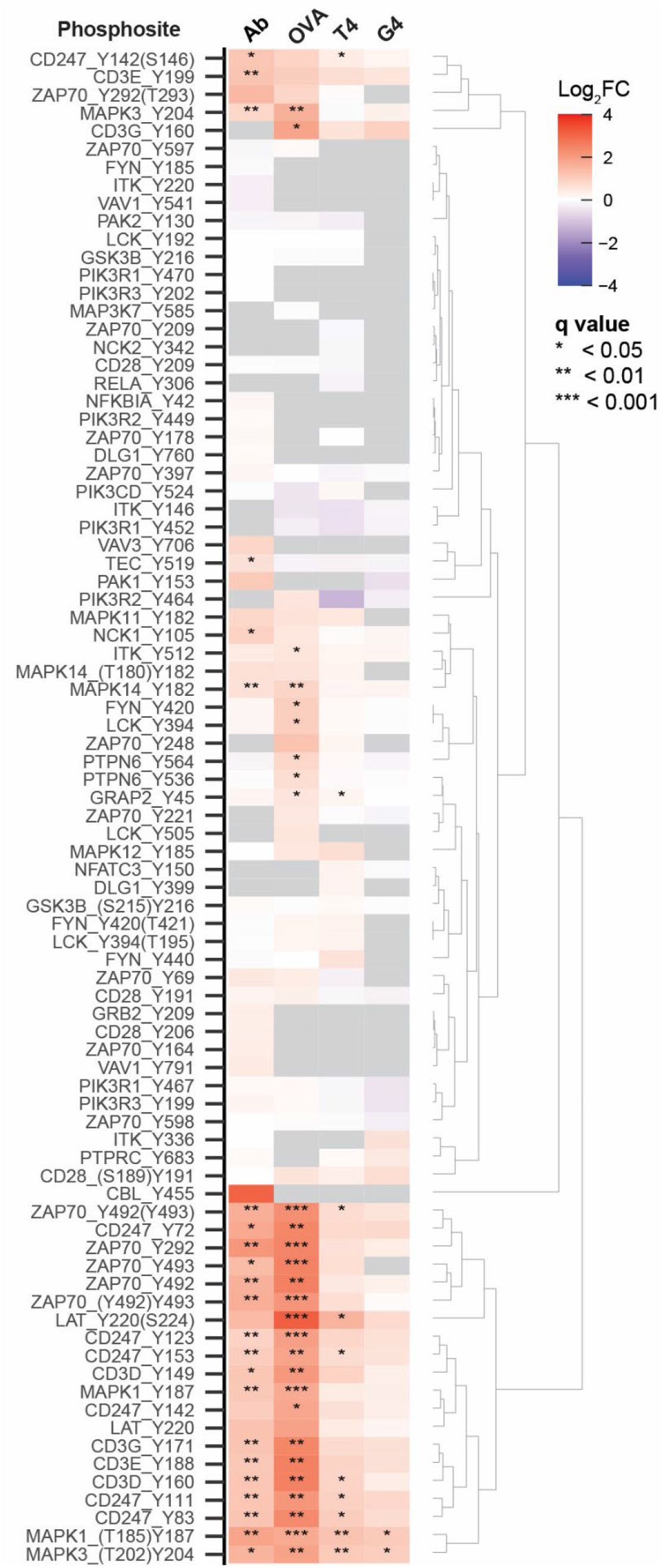
Fold change (FC) intensity heatmap of all pTyr sites involved in the TCR signaling pathway as denoted by KEGG. Hierarchical clustering was performed based on cluster distances corresponding to the dendrogram shown on the right. Each row represents a unique pTyr site following the notation [gene_name]_[Y][residue_number]. pTyr sites originating from doubly phosphorylated peptides are annotated with a parenthesis containing the second phosphorylation site with the highest localization probability. pTyr sites without a parenthesis originate from singly phosphorylated peptides.

## Discussion

In this study, we demonstrated the utility of BOOST in gaining new biological insights by comparing pTyr-mediated signaling pathways triggered by antigen-independent Ab stimulation and antigen-specific activation of Jurkat OT-1 TCR using a panel of pMHC tetramers (OVA, T4 and G4). Altered peptide ligands G4 and T4 were selected because low-affinity G4 antigen was shown to induce positive selection while T4 was reported to be the threshold affinity antigen for inducing negative selection by compartmentalizing MAPK signaling in mice thymocytes^23^. The inclusion of PV boost channel resulted in global elevation in the abundance of multiplexed pTyr-containing peptides (Figure S3A-D), enabling selective triggering of this class of precursor ions for fragmentation by DDA to facilitate reporter ion quantitation in the samples-of-interest. This strategy allows us to reproducibly quantify close to a thousand pTyr sites collectively (Figure 2F) for pathway analyses. Because the samples were enriched for pTyr-containing peptides prior to data acquisition on the mass spectrometer, it is likely more appropriate to perform pathway analyses using a PTM-centric signature database, such as PTMsigDB^38^, rather than applying a gene-centric approach like Gene Set Enrichment Analysis (GSEA). PTMsigDB provides curated phosphorylation signatures of perturbations, kinases, and signaling pathways, which is the basis for site-specific PTM-SEA deployed in this study. Interestingly, PTM-SEA reveals a close resemblance in enriched signatures between antigen-independent and antigen-specific activation of TCR signaling across all conditions, as no divergence in signaling pathways was particularly evident. Specifically, perturbation signatures that were commonly enriched by Ab, OVA, T4 and G4 such as anti-CD3, vanadate and thrombin (Figure 5A-C) were previously validated in TCR-targeting antibody-based studies^50–54^, ^58^. Altogether, our data indicate that TCR activation triggered by antigen-independent Ab or antigen-specific OVA pMHC may share many common pTyr-mediated signaling components.

In addition to the resemblance in signaling pathways between Ab and antigenic pMHC, we also observed indirect evidence that possibly indicates the initiation of a negative feedback regulation upon strong TCR activation by high-affinity antigen OVA, evidenced by several known negative regulators of TCR signaling that were differentially phosphorylated, including PTPN6, PAG1 and UBASH3A (Figure 2A,C, Figure 7). Firstly, PTPN6 (also known as SHP-1 tyrosine phosphatase) has been shown to dephosphorylate the activation site of the initiating kinase Lck Y394 to destabilize its active conformation and promote the inactive closed confirmation as a mechanism to suppress T cell signaling^60^. Notably, two tyrosine sites in the C-terminus of PTPN6 (Y536 and Y564) that were significantly upregulated by OVA activation in our data (Figure 2C, Figure 7) have been shown to correlate with increased phosphatase activity upon the phosphorylation of those sites^59^. It was hypothesized that PTPN6 promotes negative feedback of TCR signaling by inhibiting/competing with the positive feedback regulation by MAPK^61^. Intriguingly, the dual phosphorylation activation sites of MAPK1 and MAPK3 of OVA were ~2-fold higher than T4 or G4 in our data (Figure 4). Secondly, PAG1 was proposed as a negative regulator of TCR signaling by recruiting Csk to phosphorylate the inhibitory tyrosine site of Lck^62^. A quantitative phosphoproteomic study reported a negative feedback loop mediated by SLP-76 by regulating the phosphorylation of PAG1 Y227^63^, and this pTyr site was significantly increased in OVA-triggered activation of TCR (Figure 2C). Thirdly, UBASH3A has been previously reported to associate with E3 ubiquitin ligase CBL and suppresses TCR signaling via a ubiquitin-dependent mechanism^64, 65^, and we observed a significant upregulation of UBASH3A Y3 phosphosite in the OVA-activated sample. Interestingly, we recently discovered UBASH3A as one of the proximal Lck interactors upon TCR activation using Lck-TurboID proximity labeling^66^. Both sets of data are consistent with our hypothesis that Lck interacts with and phosphorylates UBASH3A Y9, or facilitates the association of another tyrosine kinase with UBASH3A to phosphorylate Y9, that enables the assembly of ubiquitination machinery in the degradation of TCR signaling complex. An extension of this study investigating the temporal effect of ubiquitination levels in high- and low-affinity antigens may help test this hypothesis. Collectively, our data suggest possible mechanisms of negative feedback regulation to dampen TCR signaling after a strong stimulation induced by a high-affinity antigen.

Apart from elucidating the mechanisms of T cell signaling pathways, phosphosite-specific pathway analyses can potentially be extended to expand our understanding in disease mechanisms. For instance, in our KEGG pathway analysis, differential pTyr sites in our data are highly represented in Chagas disease (Figure 6) caused by the parasite *Trypanosoma cruzi*. Despite after more than a century after its initial discovery, Chagas disease (also known as American trypanosomiasisis) remains an incurable tropical disease and continues to infect millions of people worldwide due to the enigmatic underlying causes of the infection^67^. Although it is known that patients of Chagas disease contain elevated levels of pro-inflammatory cytokines released by CD8+ cytotoxic T cells^67^, the underlying molecular mechanism is still incompletely understood. The BOOST approach here may provide a framework for future studies of T cells in the context of Chagas disease states. However, we acknowledge that clinical applications are only possible if more validation studies are conducted to carefully test the hypotheses generated from these proteomic data.

To maximize our data quality for meaningful interpretation of the data, we carefully optimized the BOOST methodology employed in this study by considering many aspects during pre- and post-data acquisition. First, to ensure that changes in pTyr sites abundance are stochiometric and not due to the changes in protein expression, we used the same (isogenic) cell line for all analysis to minimize variation from cellular heterogeneity between different cell lines. We also carefully performed an additional normalization step prior to TMT pooling based on the median intensity of each individually labeled peptides to ensure that the stimulated samples and controls were pooled in equal amounts. In terms of TMT channel designation, we designated TMT126 as the PV boost channel, followed by 2 blank channels in TMT127N and TMT127C (Figure 1C). Following data acquisition, we verified that reporter interference of PV boost channel resulting from 13C and 15N isotopic impurities from the labeling reagent can clearly be observed, resulting in false positives in the blank channels (Figure S3A-D). The incorporation of these blank channels helped circumvent reporter leakage into the sample channels which could negatively skew reporter ion quantitation. Not only was +2 leakage into TMT128N and TMT128C not observed, the reporter intensities of TMT128N and TMT128C were also not significantly different from their corresponding replicates (Figure S3A-D), suggesting that reporter interference from PV boost channel was limited to +1 leakage into TMT127N and TMT127C only. Although reporter interference can similarly be observed in non-pTyr PSMs, it is more pronounced in pTyr-containing PSMs (Figure S3A-D). It is possible that the placement of blank channels minimized quantitative contortion in the samples-of-interest, resulting in median CVs of less than 20% in pTyr PSM across all conditions (Figure S4). This suggests that precision and ion sampling in the quantitation of reporter ions was reasonably good as benchmarked by a single-cell proteomics study utilizing a carrier channel where ≤20% CV was defined as accurate quantification^68^.

We attribute the observed high quantitative precision to a median boost level of 4-to 30-fold specific to pTyr-containing PSMs only (Figure S5). Expectedly, the median boost levels positively correlate with the number of missing values within the reporter channels per pTyr PSM (Figure S5). This happens because we hypothesize that PV ions tend to be disproportionately overrepresented in the Orbitrap mass analyzer during the sampling of low-abundance sample ions, while low-abundance ions are more likely to result in missing values due to certain signal-to-noise threshold imposed by quantitation software such as MaxQuant^69^. Furthermore, this effect of boost level is specific to pTyr PSM only (Figure S5). A boost level of 4-to 30-fold is in close agreement with the recommendation of a recent single-cell proteomics study suggesting a carrier boost channel of ~20X for optimal ion sampling and minimal ion coalescence or space charging effects in Orbitrap instruments^68^. A recent evaluation of BOOST reported that PV boost channel adversely impacted the quantitative accuracy of reporter ions due to severe ratio compression, high CV% (up to median 78%), and isotopic interference^70^ using a MS2-based TMT quantitation approach. However, MS2-based TMT quantitation is known to suffer from ratio compression^71^, and we have previously shown that contrived ratios closely matched expected values in our SPS-MS3-based quantitation^6^. Here, we minimized the impact of isotopic interference from PV boost channel by strategic placement of PV boost channel in TMT126 with TMT127N and TMT127C left empty. We further validated our quantitation from proteomic data by comparing directly with conventional immunoblot quantitation and found consistent agreement among both methods (Figure 4). We attempted to validate other pTyr sites (such as Zap70 Y493 and LAT Y220) using other commercially available antibodies but these antibodies could only detect PV-treated samples containing highly abundant pTyr sites at non-physiological levels (Figure S6). This underscores the benefits of BOOST in overcoming the limitations of using phospho-specific antibodies to study phosphorylation-driven cellular signaling. We believe the optimized BOOST design shown in this study minimized technical artefacts that might negatively impact quantitation, providing confidence in the interpretation of the biological insights of the data.

In conclusion, our data suggest that pTyr-mediated regulatory axis triggered by OVA antigen-specific activation of TCR closely resembled that of antigen-independent stimulation using anti-TCR antibody, albeit OVA likely induced a relatively stronger signaling effect. While data from this study do not invalidate previous studies of T cell signaling using antibody-based stimulation, our data revealed potential advantages of using pMHC tetramers in studying T cell signaling. Antigen-specific activation of the OT-1 TCR using a panel of pMHC tetramers (OVA, T4 and G4) generated data that correlate well with the corresponding binding affinity of the pMHC, consistent with predicted signaling strength. Importantly, the apparent correlation between signaling strength and pMHC affinity enables us to fine-tune the signaling strength of T cell stimulation in future studies, allowing a more precise control of the experimental parameter instead of a more binary antibody-based activation. The utility of BOOST can also potentially be extrapolated to other antigen-specific TCR and a wide range of binding kinetics to better understand the mechanisms of how TCR interacts with tumor-specific antigens presented by cancer cells. We acknowledge that clinical application of PTM-centric proteomics is still in its infancy, but such applications will hopefully become more feasible over time, facilitated by the curation of better PTM-centric database annotations in synergy with genomic and proteomic studies.

## SUPPORTING INFORMATION

Figure S1: Immunoblot examining the total tyrosine phosphorylation levels across the proteome. Figure S2: Histogram of the number of PSM containing at least one phosphorylation modification on serine (S), threonine (T) or tyrosine (Y) residues plotted against the localization probability of the phosphosite.

Figure S3: The total number of quantifiable unique pTyr-containing (A) and non-pTyr-containing (B) PSM was indicated for each TMT channel, including blanks (false positives). The distribution of log10 reporter ion intensities of unique pTyr-containing (C) and non-pTyr-containing (D) PSM for each TMT channel are shown in boxplots.

Figure S4: Boxplots illustrating the distribution of coefficients of variation percent (CV%) of the reporter ion intensities of unique pTyr-containing (left) and non-pTyr-containing (right) PSM for each condition.

Figure S5: Median boost levels of all unique pTyr-containing and non-pTyr-containing PSM are plotted as a function of missing values in all 8 possible reporter ion channels for each condition.

Figure S6: Immunoblot examining phosphosites using phosphosite-specific antibodies.

File S1: mqpar.xml (MaxQuant)

File S2: Interactive Heatmap

File S3-S6: R scripts

File S7: Immunoblot quantitation

Table S1: evidence.txt (MaxQuant)

Table S2: Phospho (STY)Sites.txt (MaxQuant)

Table S3: KEGG-annotated pTyr site quantitation

## Acknowledgements

The authors wish to thank Dr. Ricky Edmondson and Dr. Samuel G. Mackintosh from University of Arkansas for Medical Sciences (UAMS) for running the proteomic samples, Dr. Aaron Storey from UAMS for kindly sharing the R scripts used in PTM-SEA analysis, Dr. Arthur Weiss for kindly providing C305 antibody, and Dr. Ondrej Stepanek for generating the J.OT1 cell line used in this study. All pMHC tetramers used in this study (OVA, T4, G4 and VSV; task order 46336, 46338, 46339, 46340, respectively) were obtained through the NIH Tetramer Core Facility (Atlanta, GA). We acknowledge financial support from NIH grants R01AI083636, P01AI091580, and P20GM121293.

## Data Availability

The mass spectrometry proteomic data have been deposited to the ProteomeXchange Consortium (http://proteomecentral.proteomexchange.org) via the PRIDE partner repository ^72^ with the dataset identifier PXD024579.

## References

1. Abraham, R. T.; Weiss, A., Jurkat T cells and development of the T-cell receptor signalling paradigm. Nat Rev Immunol 2004, 4 (4), 301–8.

2. Love, P. E.; Hayes, S. M., ITAM-mediated signaling by the T-cell antigen receptor. Cold Spring Harb Perspect Biol 2010, 2 (6), a002485.

3. Hunter, T., Protein modification: phosphorylation on tyrosine residues. Curr Opin Cell Biol 1989, 1 (6), 1168–81.

4. Bian, Y.; Li, L.; Dong, M.; Liu, X.; Kaneko, T.; Cheng, K.; Liu, H.; Voss, C.; Cao, X.; Wang, Y.; Litchfield, D.; Ye, M.; Li, S. S.; Zou, H., Ultra-deep tyrosine phosphoproteomics enabled by a phosphotyrosine superbinder. Nat Chem Biol 2016, 12 (11), 959–966.

5. Mandell, J. W., Phosphorylation state-specific antibodies: applications in investigative and diagnostic pathology. Am J Pathol 2003, 163 (5), 1687–98.

6. Chua, X. Y.; Mensah, T.; Aballo, T.; Mackintosh, S. G.; Edmondson, R. D.; Salomon, A. R., Tandem Mass Tag Approach Utilizing Pervanadate BOOST Channels Delivers Deeper Quantitative Characterization of the Tyrosine Phosphoproteome. Molecular & cellular proteomics: MCP 2020, 19 (4), 730–743.

7. Huyer, G.; Liu, S.; Kelly, J.; Moffat, J.; Payette, P.; Kennedy, B.; Tsaprailis, G.; Gresser, M. J.; Ramachandran, C., Mechanism of inhibition of protein-tyrosine phosphatases by vanadate and pervanadate. The Journal of biological chemistry 1997, 272 (2), 843–51.

8. de Wet, B.; Zech, T.; Salek, M.; Acuto, O.; Harder, T., Proteomic characterization of plasma membrane-proximal T cell activation responses. The Journal of biological chemistry 2011, 286 (6), 4072–80.

9. Lichtenfels, R.; Rappl, G.; Hombach, A. A.; Recktenwald, C. V.; Dressler, S. P.; Abken, H.; Seliger, B., A proteomic view at T cell costimulation. PloS one 2012, 7 (4), e32994.

10. Weiss, A.; Stobo, J. D., Requirement for the coexpression of T3 and the T cell antigen receptor on a malignant human T cell line. J Exp Med 1984, 160 (5), 1284–99.

11. Van Wauwe, J. P.; De Mey, J. R.; Goossens, J. G., OKT3: a monoclonal anti-human T lymphocyte antibody with potent mitogenic properties. Journal of immunology (Baltimore, Md.: 1950) 1980, 124 (6), 2708–13.

12. Lo, Y. C.; Edidin, M. A.; Powell, J. D., Selective activation of antigen-experienced T cells by anti-CD3 constrained on nanoparticles. Journal of immunology (Baltimore, Md.: 1950) 2013, 191 (10), 5107–14.

13. Poltorak, M. P.; Graef, P.; Tschulik, C.; Wagner, M.; Cletiu, V.; Dreher, S.; Borjan, B.; Fraessle, S. P.; Effenberger, M.; Turk, M.; Busch, D. H.; Plitzko, J.; Kugler, D. G.; Ragan, S.; Schmidt, T.; Stemberger, C.; Germeroth, L., Expamers: a new technology to control T cell activation. Sci Rep 2020, 10 (1), 17832.

14. Jacobs, N.; Mazzoni, A.; Mezzanzanica, D.; Negri, D. R.; Valota, O.; Colnaghi, M. I.; Moutschen, M. P.; Boniver, J.; Canevari, S., Efficiency of T cell triggering by anti-CD3 monoclonal antibodies (mAb) with potential usefulness in bispecific mAb generation. Cancer Immunol Immunother 1997, 44 (5), 257–64.

15. Li, Y.; Kurlander, R. J., Comparison of anti-CD3 and anti-CD28-coated beads with soluble anti-CD3 for expanding human T cells: differing impact on CD8 T cell phenotype and responsiveness to restimulation. J Transl Med 2010, 8, 104.

16. Kelly, J. M.; Sterry, S. J.; Cose, S.; Turner, S. J.; Fecondo, J.; Rodda, S.; Fink, P. J.; Carbone, F. R., Identification of conserved T cell receptor CDR3 residues contacting known exposed peptide side chains from a major histocompatibility complex class I-bound determinant. Eur J Immunol 1993, 23 (12), 3318–26.

17. Clarke, S. R.; Barnden, M.; Kurts, C.; Carbone, F. R.; Miller, J. F.; Heath, W. R., Characterization of the ovalbumin-specific TCR transgenic line OT-I: MHC elements for positive and negative selection. Immunol Cell Biol 2000, 78 (2), 110–7.

18. Gascoigne, N. R.; Rybakin, V.; Acuto, O.; Brzostek, J., TCR Signal Strength and T Cell Development. Annu Rev Cell Dev Biol 2016, 32, 327–348.

19. Rotzschke, O.; Falk, K.; Stevanovic, S.; Jung, G.; Walden, P.; Rammensee, H. G., Exact prediction of a natural T cell epitope. Eur J Immunol 1991, 21 (11), 2891–4.

20. Alam, S. M.; Davies, G. M.; Lin, C. M.; Zal, T.; Nasholds, W.; Jameson, S. C.; Hogquist, K. A.; Gascoigne, N. R.; Travers, P. J., Qualitative and quantitative differences in T cell receptor binding of agonist and antagonist ligands. Immunity 1999, 10 (2), 227–37.

21. Lo, W. L.; Shah, N. H.; Ahsan, N.; Horkova, V.; Stepanek, O.; Salomon, A. R.; Kuriyan, J.; Weiss, A., Lck promotes Zap70-dependent LAT phosphorylation by bridging Zap70 to LAT. Nat Immunol 2018, 19 (7), 733–741.

22. Hogquist, K. A.; Jameson, S. C.; Heath, W. R.; Howard, J. L.; Bevan, M. J.; Carbone, F. R., T cell receptor antagonist peptides induce positive selection. Cell 1994, 76 (1), 17–27.

23. Daniels, M. A.; Teixeiro, E.; Gill, J.; Hausmann, B.; Roubaty, D.; Holmberg, K.; Werlen, G.; Hollander, G. A.; Gascoigne, N. R.; Palmer, E., Thymic selection threshold defined by compartmentalization of Ras/MAPK signalling. Nature 2006, 444 (7120), 724–9.

24. Chakraborty, A. K.; Weiss, A., Insights into the initiation of TCR signaling. Nat Immunol 2014, 15 (9), 798–807.

25. Rosette, C.; Werlen, G.; Daniels, M. A.; Holman, P. O.; Alam, S. M.; Travers, P. J.; Gascoigne, N. R.; Palmer, E.; Jameson, S. C., The impact of duration versus extent of TCR occupancy on T cell activation: a revision of the kinetic proofreading model. Immunity 2001, 15 (1), 59–70.

26. Martin, A.; Tisch, R. M.; Getts, D. R., Manipulating T cell-mediated pathology: targets and functions of monoclonal antibody immunotherapy. Clin Immunol 2013, 148 (1), 136–47.

27. Bianchi, V.; Harari, A.; Coukos, G., Neoantigen-Specific Adoptive Cell Therapies for Cancer: Making T-Cell Products More Personal. Front Immunol 2020, 11, 1215.

28. Rassy, E.; Flippot, R.; Albiges, L., Tyrosine kinase inhibitors and immunotherapy combinations in renal cell carcinoma. Ther Adv Med Oncol 2020, 12, 1758835920907504.

29. Altman, J. D.; Moss, P. A.; Goulder, P. J.; Barouch, D. H.; McHeyzer-Williams, M. G.; Bell, J. I.; McMichael, A. J.; Davis, M. M., Phenotypic analysis of antigen-specific T lymphocytes. Science (New York, N.Y.) 1996, 274 (5284), 94–6.

30. Wisniewski, J. R.; Zougman, A.; Nagaraj, N.; Mann, M., Universal sample preparation method for proteome analysis. Nat Methods 2009, 6 (5), 359–62.

31. Ahsan, N.; Salomon, A. R., Quantitative Phosphoproteomic Analysis of T-Cell Receptor Signaling. Methods in molecular biology (Clifton, N.J.) 2017, 1584, 369–382.

32. Brenes, A.; Hukelmann, J. L.; Bensaddek, D.; Lamond, A. I., Multi-batch TMT reveals false positives, batch effects and missing values. Molecular & cellular proteomics: MCP 2019.

33. McAlister, G. C.; Nusinow, D. P.; Jedrychowski, M. P.; Wuhr, M.; Huttlin, E. L.; Erickson, B. K.; Rad, R.; Haas, W.; Gygi, S. P., MultiNotch MS3 enables accurate, sensitive, and multiplexed detection of differential expression across cancer cell line proteomes. Analytical chemistry 2014, 86 (14), 7150–8.

34. Cox, J.; Mann, M., MaxQuant enables high peptide identification rates, individualized p.p.b.-range mass accuracies and proteome-wide protein quantification. Nat Biotechnol 2008, 26 (12), 1367–72.

35. Cox, J.; Neuhauser, N.; Michalski, A.; Scheltema, R. A.; Olsen, J. V.; Mann, M., Andromeda: a peptide search engine integrated into the MaxQuant environment. Journal of proteome research 2011, 10 (4), 1794–805.

36. Storey, J. D.; Tibshirani, R., Statistical significance for genomewide studies. Proc Natl Acad Sci U S A 2003, 100 (16), 9440–5.

37. Tyanova, S.; Temu, T.; Sinitcyn, P.; Carlson, A.; Hein, M. Y.; Geiger, T.; Mann, M.; Cox, J., The Perseus computational platform for comprehensive analysis of (prote)omics data. Nat Methods 2016, 13 (9), 731–40.

38. Krug, K.; Mertins, P.; Zhang, B.; Hornbeck, P.; Raju, R.; Ahmad, R.; Szucs, M.; Mundt, F.; Forestier, D.; Jane-Valbuena, J.; Keshishian, H.; Gillette, M. A.; Tamayo, P.; Mesirov, J. P.; Jaffe, J. D.; Carr, S.; Mani, D. R., A Curated Resource for Phosphosite-specific Signature Analysis. Molecular & cellular proteomics: MCP 2019, 18 (3), 576–593.

39. Storey, A. J.; Naceanceno, K. S.; Lan, R. S.; Washam, C. L.; Orr, L. M.; Mackintosh, S. G.; Tackett, A. J.; Edmondson, R. D.; Wang, Z.; Li, H. Y.; Frett, B.; Kendrick, S.; Byrum, S. D., ProteoViz: a tool for the analysis and interactive visualization of phosphoproteomics data. Mol Omics 2020.

40. Hardwick, J. S.; Sefton, B. M., The activated form of the Lck tyrosine protein kinase in cells exposed to hydrogen peroxide is phosphorylated at both Tyr-394 and Tyr-505. The Journal of biological chemistry 1997, 272 (41), 25429–32.

41. Chan, A. C.; Dalton, M.; Johnson, R.; Kong, G. H.; Wang, T.; Thoma, R.; Kurosaki, T., Activation of ZAP-70 kinase activity by phosphorylation of tyrosine 493 is required for lymphocyte antigen receptor function. EMBO J 1995, 14 (11), 2499–508.

42. Paz, P. E.; Wang, S.; Clarke, H.; Lu, X.; Stokoe, D.; Abo, A., Mapping the Zap-70 phosphorylation sites on LAT (linker for activation of T cells) required for recruitment and activation of signalling proteins in T cells. Biochem J 2001, 356 (Pt 2), 461–71.

43. Heyeck, S. D.; Wilcox, H. M.; Bunnell, S. C.; Berg, L. J., Lck phosphorylates the activation loop tyrosine of the Itk kinase domain and activates Itk kinase activity. The Journal of biological chemistry 1997, 272 (40), 25401–8.

44. Sekiya, F.; Poulin, B.; Kim, Y. J.; Rhee, S. G., Mechanism of tyrosine phosphorylation and activation of phospholipase C-gamma 1. Tyrosine 783 phosphorylation is not sufficient for lipase activation. The Journal of biological chemistry 2004, 279 (31), 32181–90.

45. Kortum, R. L.; Rouquette-Jazdanian, A. K.; Samelson, L. E., Ras and extracellular signal-regulated kinase signaling in thymocytes and T cells. Trends Immunol 2013, 34 (6), 259–68.

46. Morrison, D. K., MAP kinase pathways. Cold Spring Harb Perspect Biol 2012, 4 (11).

47. Carreno, M.; Fuller, L.; Zucker, K.; Yang, W. C.; Burke, G.; Nery, J.; Gomez, C.; Esquenazi, V.; Miller, J., Cross-species reactivity of the anti-idiotype anti-OKT3 cascade between mice and humans. Hum Immunol 1992, 33 (4), 249–58.

48. Kuhn, C.; Weiner, H. L., Therapeutic anti-CD3 monoclonal antibodies: from bench to bedside. Immunotherapy 2016, 8 (8), 889–906.

49. Biffen, M.; Shiroo, M.; Alexander, D. R., G-proteins are not directly involved in the CD3-antigen-mediated production of inositol phosphates in HPB-ALL T-leukaemia cells expressing phospholipase C isoforms gamma 1 and beta 3. Biochem J 1993, 289 (Pt 2), 387–94.

50. Deira, J.; Alberca, I.; Lerma, J. L.; Martin, B.; Tabernero, J. M., Changes in coagulation and fibrinolysis in the postoperative period immediately after kidney transplantation in patients receiving OKT3 or cyclosporine A as induction therapy. Am J Kidney Dis 1998, 32 (4), 575–81.

51. Valmu, L.; Gahmberg, C. G., Treatment with okadaic acid reveals strong threonine phosphorylation of CD18 after activation of CD11/CD18 leukocyte integrins with phorbol esters or CD3 antibodies. Journal of immunology (Baltimore, Md.: 1950) 1995, 155 (3), 1175–83.

52. Poujol, F.; Monneret, G.; Pachot, A.; Textoris, J.; Venet, F., Altered T Lymphocyte Proliferation upon Lipopolysaccharide Challenge Ex Vivo. PloS one 2015, 10 (12), e0144375.

53. Sherry, N.; Hagopian, W.; Ludvigsson, J.; Jain, S. M.; Wahlen, J.; Ferry, R. J., Jr.; Bode, B.; Aronoff, S.; Holland, C.; Carlin, D.; King, K. L.; Wilder, R. L.; Pillemer, S.; Bonvini, E.; Johnson, S.; Stein, K. E.; Koenig, S.; Herold, K. C.; Daifotis, A. G.; Protege Trial, I., Teplizumab for treatment of type 1 diabetes (Protege study): 1-year results from a randomised, placebo-controlled trial. Lancet 2011, 378 (9790), 487–97.

54. Yiemwattana, I.; Ngoenkam, J.; Paensuwan, P.; Kriangkrai, R.; Chuenjitkuntaworn, B.; Pongcharoen, S., Essential role of the adaptor protein Nck1 in Jurkat T cell activation and function. Clin Exp Immunol 2012, 167 (1), 99–107.

55. Lindauer, M.; Hochhaus, A., Dasatinib. Recent Results Cancer Res 2018, 212, 29–68.

56. Courtney, A. H.; Lo, W. L.; Weiss, A., TCR Signaling: Mechanisms of Initiation and Propagation. Trends Biochem Sci 2018, 43 (2), 108–123.

57. Krumenacker, J. S.; Montgomery, D. W.; Buckley, D. J.; Gout, P. W.; Buckley, A. R., Prolactin receptor signaling: shared components with the T-cell antigen receptor in Nb2 lymphoma cells. Endocrine 1998, 9 (3), 313–20.

58. Montgomery, D. W.; Krumenacker, J. S.; Buckley, A. R., Prolactin stimulates phosphorylation of the human T-cell antigen receptor complex and ZAP-70 tyrosine kinase: a potential mechanism for its immunomodulation. Endocrinology 1998, 139 (2), 811–4.

59. Lorenz, U., SHP-1 and SHP-2 in T cells: two phosphatases functioning at many levels. Immunol Rev 2009, 228 (1), 342–59.

60. Chiang, G. G.; Sefton, B. M., Specific dephosphorylation of the Lck tyrosine protein kinase at Tyr-394 by the SHP-1 protein-tyrosine phosphatase. The Journal of biological chemistry 2001, 276 (25), 23173–8.

61. Stefanova, I.; Hemmer, B.; Vergelli, M.; Martin, R.; Biddison, W. E.; Germain, R. N., TCR ligand discrimination is enforced by competing ERK positive and SHP-1 negative feedback pathways. Nat Immunol 2003, 4 (3), 248–54.

62. Hrdinka, M.; Horejsi, V., PAG--a multipurpose transmembrane adaptor protein. Oncogene 2014, 33 (41), 4881–92.

63. Cao, L.; Ding, Y.; Hung, N.; Yu, K.; Ritz, A.; Raphael, B. J.; Salomon, A. R., Quantitative phosphoproteomics reveals SLP-76 dependent regulation of PAG and Src family kinases in T cells. PloS one 2012, 7 (10), e46725.

64. Ge, Y.; Paisie, T. K.; Newman, J. R. B.; McIntyre, L. M.; Concannon, P., UBASH3A Mediates Risk for Type 1 Diabetes Through Inhibition of T-Cell Receptor-Induced NF-kappaB Signaling. Diabetes 2017, 66 (7), 2033–2043.

65. Ge, Y.; Paisie, T. K.; Chen, S.; Concannon, P., UBASH3A Regulates the Synthesis and Dynamics of TCR-CD3 Complexes. Journal of immunology (Baltimore, Md.: 1950) 2019, 203 (11), 2827–2836.

66. Chua, X. Y.; Aballo, T.; Elnemer, W.; Tran, M.; Salomon, A., Quantitative Interactomics of Lck-TurboID in Living Human T Cells Unveils T Cell Receptor Stimulation-Induced Proximal Lck Interactors. Journal of proteome research 2021, 20 (1), 715–726.

67. Acevedo, G. R.; Girard, M. C.; Gomez, K. A., The Unsolved Jigsaw Puzzle of the Immune Response in Chagas Disease. Front Immunol 2018, 9, 1929.

68. Cheung, T. K.; Lee, C. Y.; Bayer, F. P.; McCoy, A.; Kuster, B.; Rose, C. M., Defining the carrier proteome limit for single-cell proteomics. Nat Methods 2021, 18 (1), 76–83.

69. Tyanova, S.; Temu, T.; Cox, J., The MaxQuant computational platform for mass spectrometry-based shotgun proteomics. Nat Protoc 2016, 11 (12), 2301–2319.

70. Stopfer, L. E.; Conage-Pough, J. E.; White, F. M., Quantitative consequences of protein carriers in immunopeptidomics and tyrosine phosphorylation MS2 analyses. bioRxiv 2021.

71. Hogrebe, A.; von Stechow, L.; Bekker-Jensen, D. B.; Weinert, B. T.; Kelstrup, C. D.; Olsen, J. V., Benchmarking common quantification strategies for large-scale phosphoproteomics. Nat Commun 2018, 9 (1), 1045.

72. Vizcaino, J. A.; Cote, R. G.; Csordas, A.; Dianes, J. A.; Fabregat, A.; Foster, J. M.; Griss, J.; Alpi, E.; Birim, M.; Contell, J.; O’Kelly, G.; Schoenegger, A.; Ovelleiro, D.; Perez-Riverol, Y.; Reisinger, F.; Rios, D.; Wang, R.; Hermjakob, H., The PRoteomics IDEntifications (PRIDE) database and associated tools: status in 2013. Nucleic Acids Res 2013, 41 (Database issue), D1063–9.

