## Supplementary Figures for "Ovalbumin antigen-specific activation of T cell receptor closely resembles soluble antibody stimulation as revealed by BOOST phosphotyrosine proteomics"

Table of Contents:

Figure S1: Immunoblot examining the total tyrosine phosphorylation levels across the proteome.

Figure S2: Histogram of the number of PSM containing at least one phosphorylation modification on serine (S), threonine (T) or tyrosine (Y) residues plotted against the localization probability of the phosphosite .

Figure S3: The total number of quantifiable unique pTyr-containing (A) and non-pTyr-containing (B) PSM was indicated for each TMT channel, including blanks (false positives). The distribution of $\log_{10}$ reporter ion intensities of unique pTyr-containing (C) and non-pTyr-containing (D) PSM for each TMT channel are shown in boxplots.

Figure S4: Histogram distribution of $\log_2$(reporter ion intensity) of all quantifiable pTyr sites within each TMT channel across all four comparisons.

Figure S5: Boxplots illustrating the distribution of coefficients of variation percent (CV%) of the reporter ion intensities of unique pTyr-containing (left) and non-pTyr-containing (right) PSM for each condition.

Figure S6: Median boost levels of all unique pTyr-containing and non-pTyr-containing PSM are plotted as a function of missing values in all 8 possible reporter ion channels for each condition.

Figure S7: Immunoblot examining phosphosites using phosphosite-specific antibodies.

### Supplementary Figures

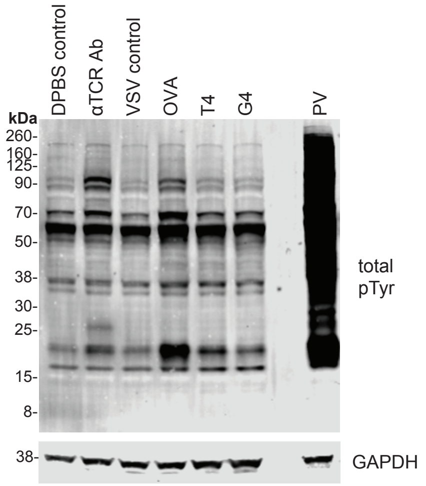


Figure S1: Immunoblot examining the total tyrosine phosphorylation levels across the proteome using an anti-phosphotyrosine antibody (4G10) of Jurkat OT1 cells treated with DPBS (no antibody control), anti-TCR antibody (Ab), and peptide-MHC loaded with VSV (null peptide), OVA, T4 and G4. GAPDH was used as a loading control.

**Figure S2**

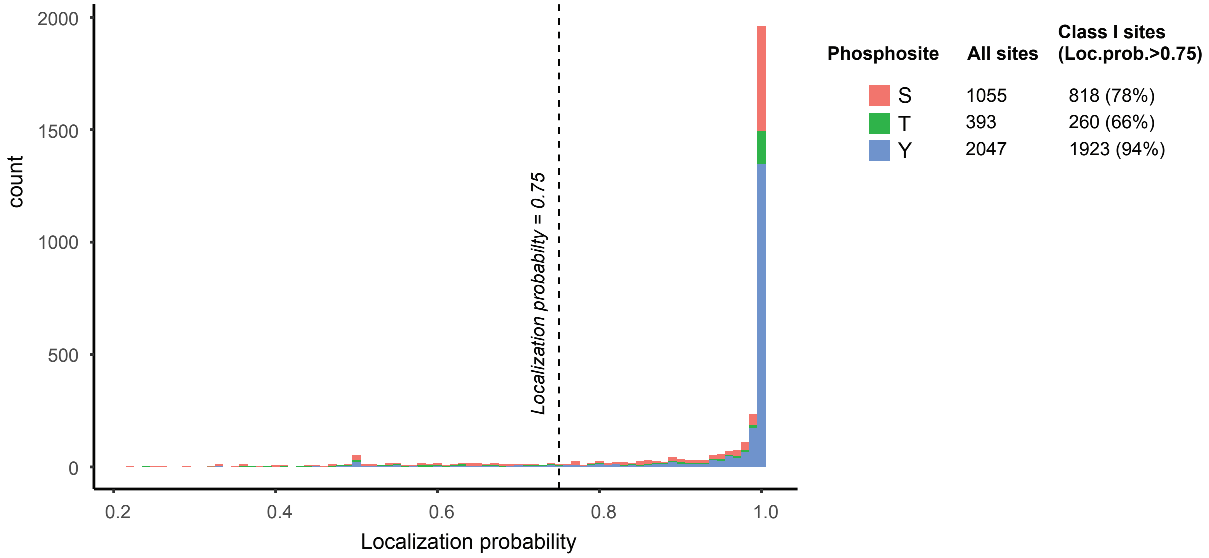


Figure S2: A histogram of the number of PSM for all conditions in the dataset containing at least one phosphorylation modification on serine (S), threonine (T) or tyrosine (Y) residues plotted against the localization probability of the phosphosite as determined by MaxQuant. The number of all sites and Class I sites (localization probability > 0.75) are indicated.

Figure S3

A

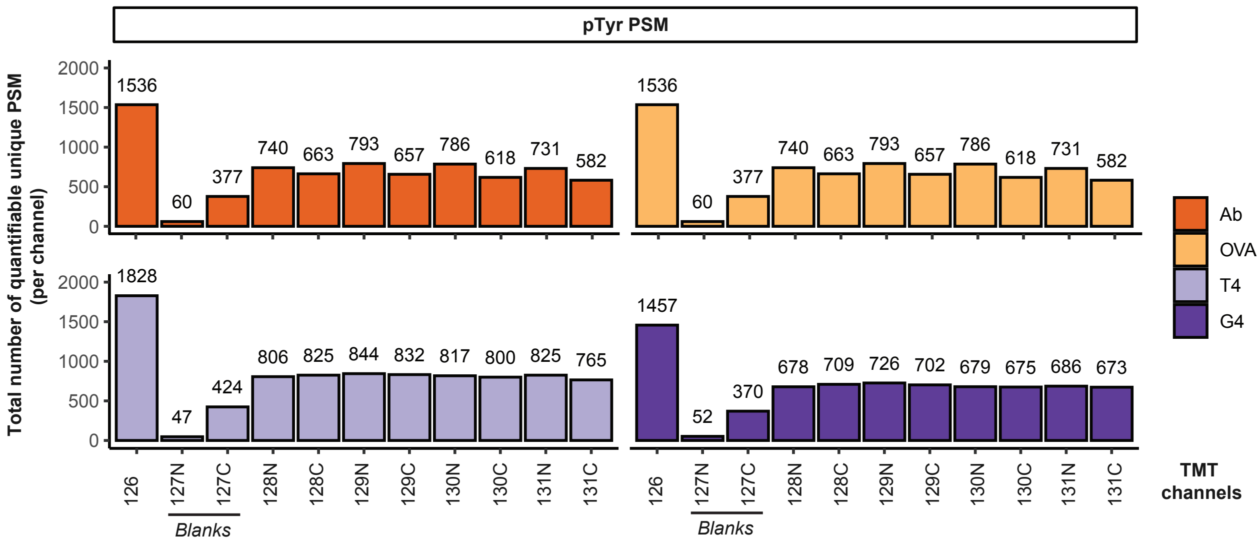
pTyr PSM
Total number of quantifiable unique PSM (per channel)
1536
60
377
740
663
793
657
786
618
731
582
1536
60
377
740
663
793
657
786
618
731
582
1828
47
424
806
825
844
832
817
800
825
765
1457
52
370
678
709
726
702
679
675
686
673
126
127N
127C
128N
128C
129N
129C
130N
130C
131N
131C
Blanks
Ab
OVA
T4
G4
TMT channels

B

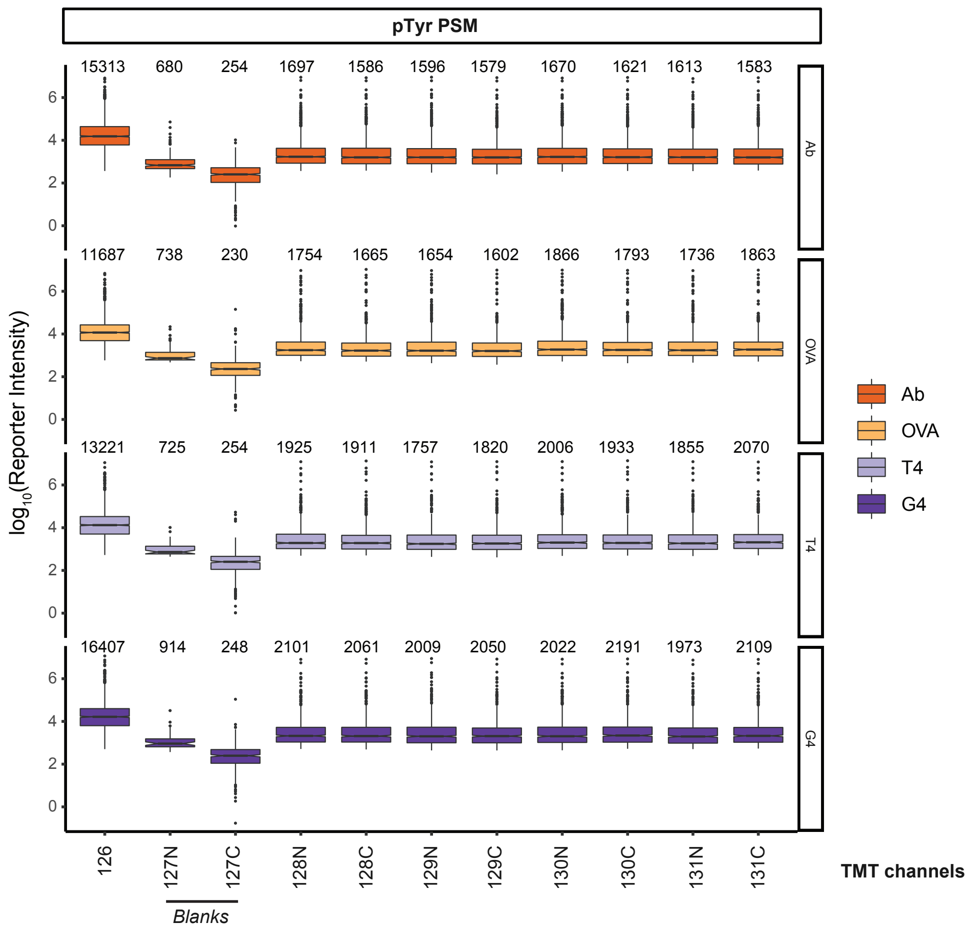
pTyr PSM
log10(Reporter Intensity)
15313 680 254 1697 1586 1596 1579 1670 1621 1613 1583
11687 738 230 1754 1665 1654 1602 1866 1793 1736 1863
13221 725 254 1925 1911 1757 1820 2006 1933 1855 2070
16407 914 248 2101 2061 2009 2050 2022 2191 1973 2109
Ab
OVA
T4
G4
126
127N
127C
128N
128C
129N
129C
130N
130C
131N
131C
Blanks
TMT channels

Figure S3

C

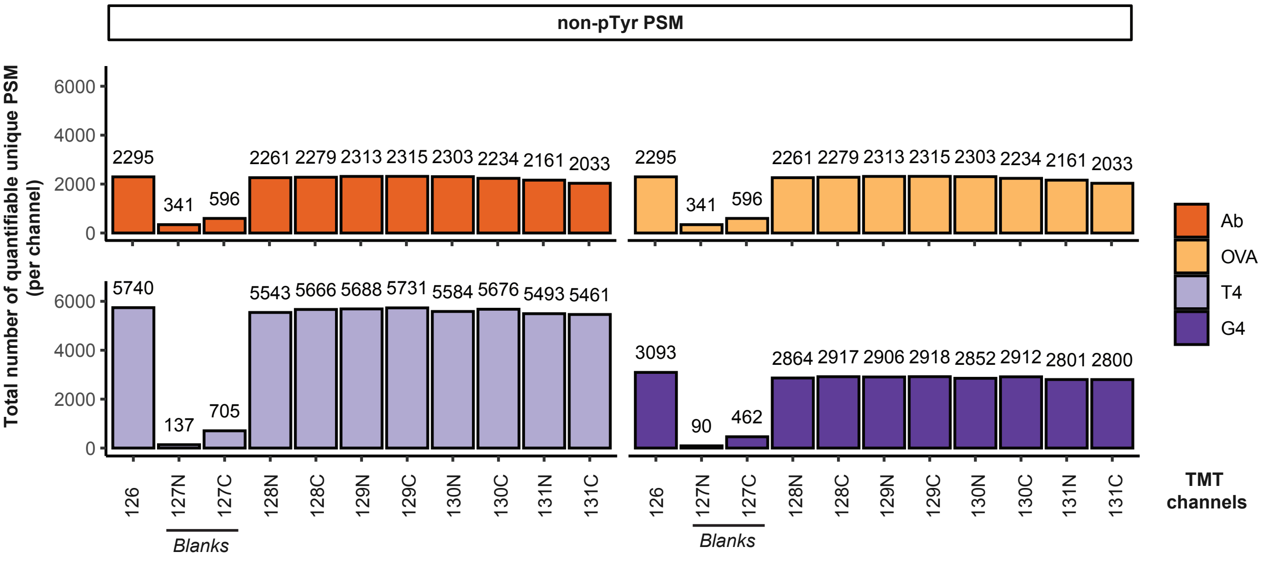
non-pTyr PSM
Total number of quantifiable unique PSM (per channel)
2295
341
596
2261 2279 2313 2315 2303 2234 2161 2033
5740
137
705
5543 5666 5688 5731 5584 5676 5493 5461
3093
90
462
2864 2917 2906 2918 2852 2912 2801 2800
Ab
OVA
T4
G4
TMT channels
126
127N
127C
128N
128C
129N
129C
130N
130C
131N
131C
Blanks

D

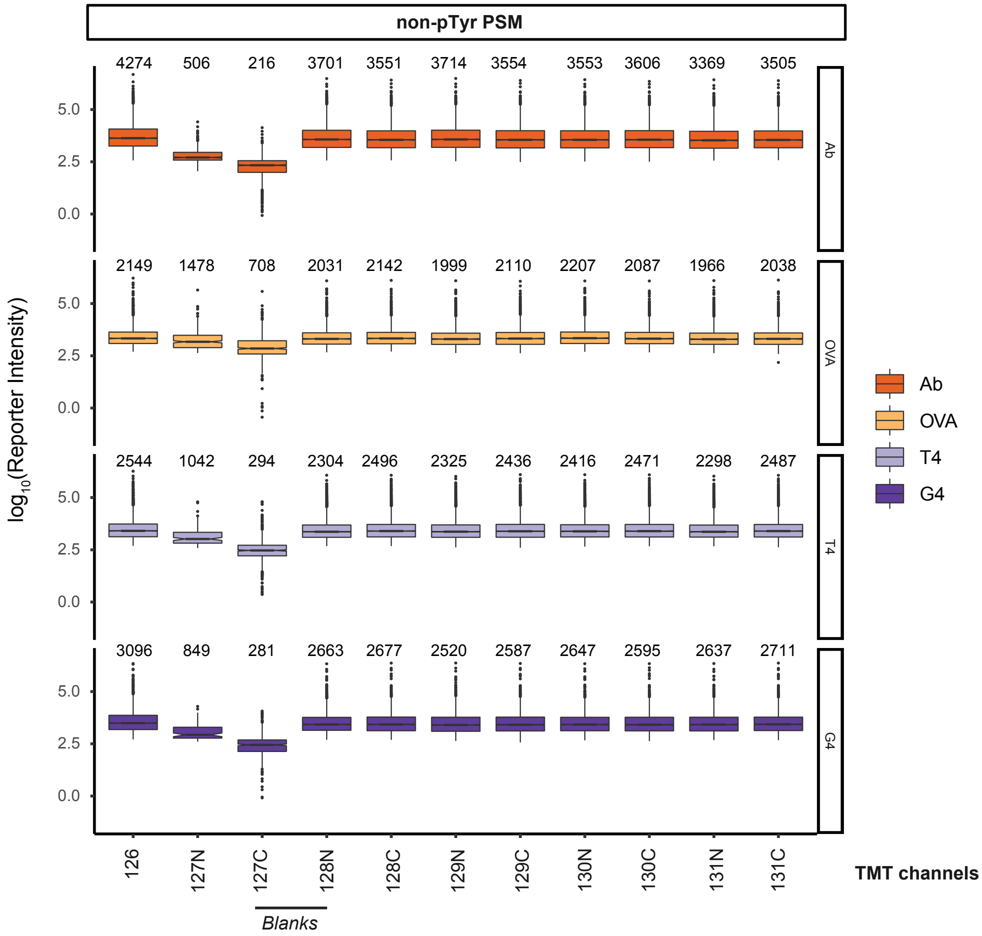
non-pTyr PSM
log10(Reporter Intensity)
4274 506 216 3701 3551 3714 3554 3553 3606 3369 3505
2149 1478 708 2031 2142 1999 2110 2207 2087 1966 2038
2544 1042 294 2304 2496 2325 2436 2416 2471 2298 2487
3096 849 281 2663 2677 2520 2587 2647 2595 2637 2711
Ab
OVA
T4
G4
TMT channels
Blanks

Figure S3: The total number of quantifiable unique pTyr-containing (A) and non-pTyr-containing (B) PSM was indicated for each TMT channel, including blanks (false positives). The distribution of $log_{10}$ reporter ion intensities of unique pTyr-containing (C) and non-pTyr-containing (D) PSM for each TMT channel are shown in boxplots. Each box represents the interquartile range of the intensities, with the middle horizontal line denoting the median of the distribution, which is also indicated above each corresponding boxplot.

**Figure S4**

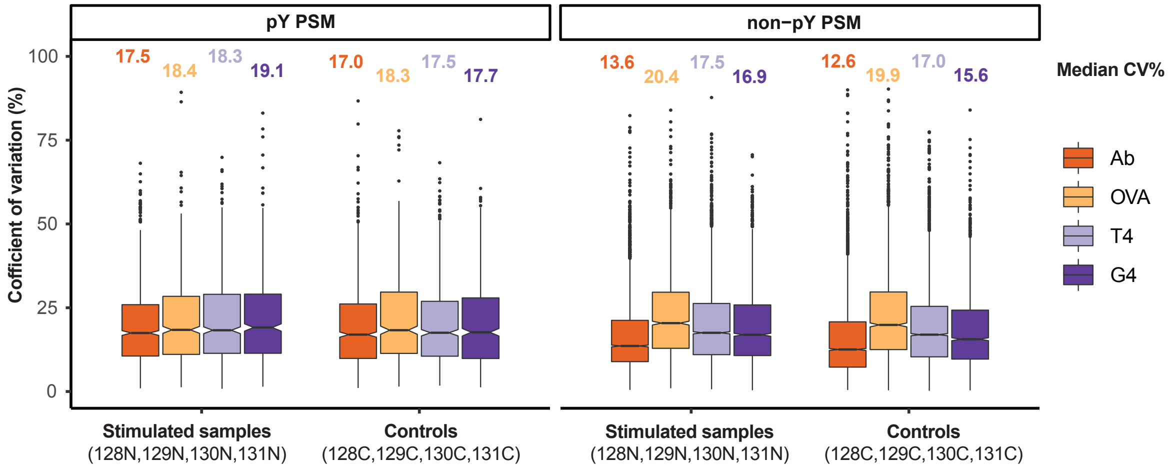


Figure S4: Boxplots illustrating the distribution of coefficients of variation percent (CV%) of the reporter ion intensities of unique pTyr-containing (left) and non-pTyr-containing (right) PSM for each condition. CV% is calculated by dividing mean by its standard deviation multiplied by 100. A minimum of 3 out of 4 possible quantified reporter ions are required for the calculation of CV%. Stimulated (Ab, OVA, T4, G4) samples originate from TMT channels 128N, 129N, 130N, and 131N, while control (DPBS, VSV, VSV, VSV) samples originate from TMT channels 128C, 129C, 130C, and 131C. Each box represents the interquartile range of the intensities, with the middle horizontal line denoting the median of the distribution, which is also indicated above each corresponding boxplot.

Figure S5

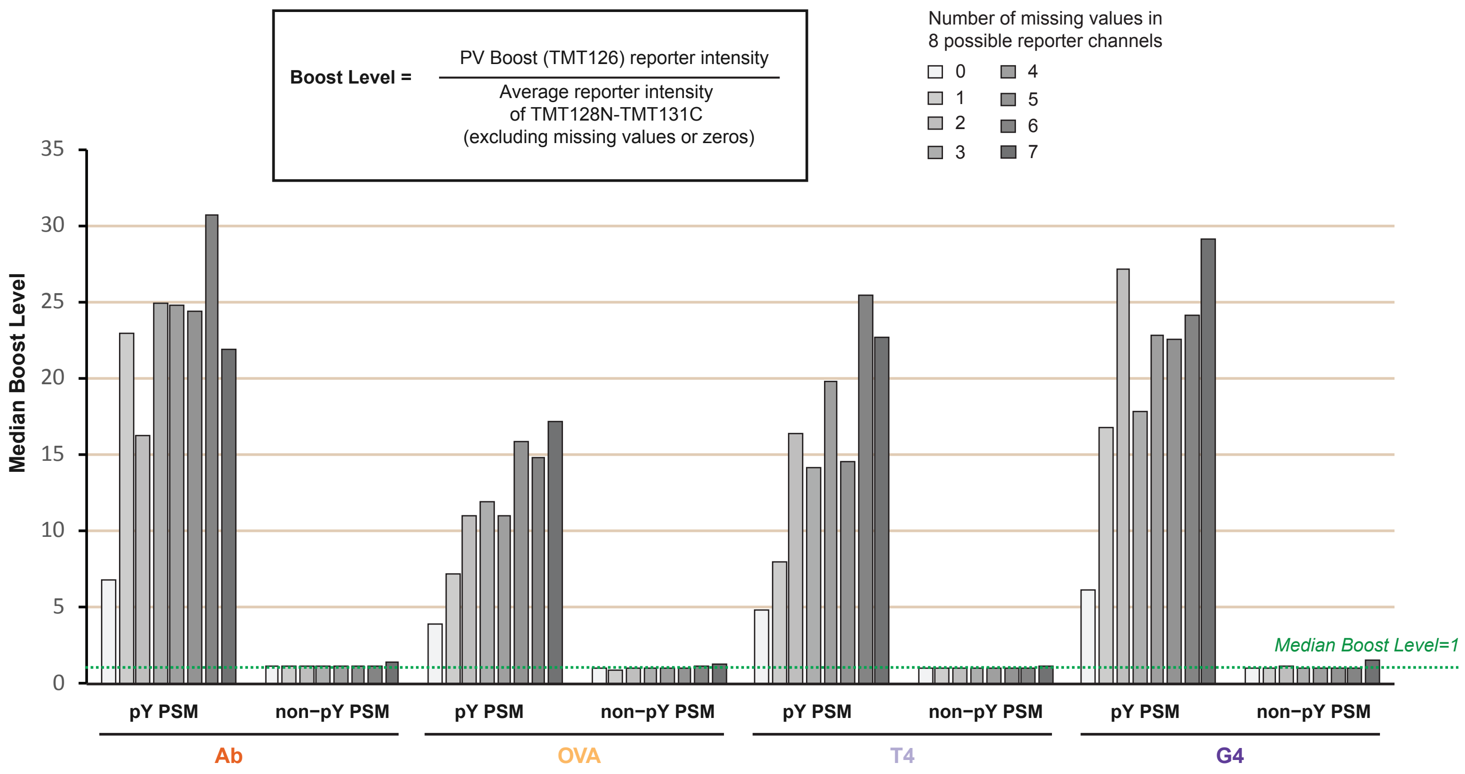


Figure S5: Median boost levels of all unique pTyr-containing and non-pTyr-containing PSM are plotted as a function of missing values in all 8 possible reporter ion channels for each condition. The calculation for boost level is shown, which is the reporter ion intensity of PV (TMT126) relative to the average of all sample reporter intensities (TMT128N to TMT131C) excluding missing values or zeros. A median boost level of 1, which indicates equivalent abundance between PV and sample for the PSM, is shown as a horizontal dotted line.

**Figure S6**

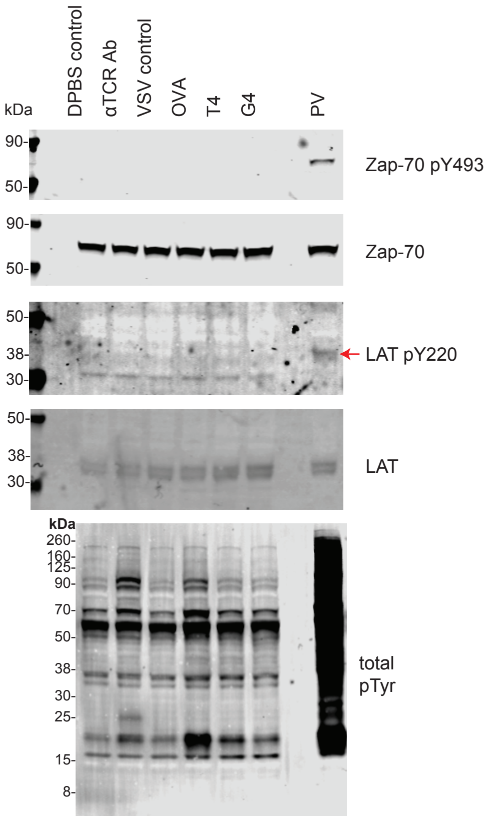


Figure S6: Immunoblot examining phosphosites of Zap-70 Y493 and LAT Y220 (indicated by a red arrow for clarity, detected in PV-treated sample only). The corresponding unphosphorylated protein of each phosphosite was used as the loading control. Total tyrosine phosphorylation level (identical to Figure S1) is shown to provide context of the effect of PV treatment.
